# Genetic analysis of grapevine wood composition identifies glycosyltransferases involved in the biosynthesis of stilbene glycosides

**DOI:** 10.64898/2026.10.24.761503

**Authors:** Hadrien Chepca, Stéphane Réty, Catarina Branco, Jean-François Chich, Geneviève Riveill, Guillaume Arnold, Gisèle Butterlin, Vincent Dumas, Marie Lahaye, Amandine Velt, Zoé Postel, Éric Duchêne, Komlan Avia, Camille Rustenholz, Raquel Tavares, Alessandra Maia-Grondard, Philippe Hugueney

**Affiliations:** Santé de la Vigne et Qualité du Vin (SVQV), INRAE, Université de Strasbourg, Colmar, France; Laboratoire de Biologie et Modélisation de la Cellule, École Normale Supérieure de Lyon, CNRS, UMR 5239, INSERM U1293, Université Claude Bernard Lyon 1, Lyon, France; Laboratoire Biométrie et Biologie Evolutive (LBBE), CNRS/Université Claude Bernard Lyon 1, Villeurbanne, France; CIBIO, Centro de Investigação em Biodiversidade e Recursos Genéticos, InBIO, Laboratório Associado, BIOPOLIS Program in Genomics, Biodiversity and Land Planning, Campus de Vairão, Departamento de Biologia, Faculdade de Ciências, Universidade do Porto, Vairão, Portugal

## Abstract

Although wood extractives may play important roles in defense against pathogens, few studies have explored the genetic determinism of their accumulation. In this work, we have used a non-targeted metabolomic approach to perform a genetic analysis of grapevine (*Vitis vinifera*) wood extractive composition in a progeny from a cross between Riesling and Gewurztraminer. Metabolic quantitative trait loci (mQTL) analyses of cane and trunk wood extractives revealed a number of mQTLs. Analysis of one of these mQTL has led to the identification of a cluster of six UDP glycosyltransferase (UGT) genes linked to the accumulation of resveratrol-glucoside (piceid) in both canes and trunks. Detailed biochemical and structural characterization of these six UGTs highlighted a surprising functional diversity among this small gene family, as well as one UGT gene as a key determinant of the constitutive accumulation of piceid in lignified grapevine tissues. Molecular evolution analyses showed that the UGTs in this gene cluster were globally under strong purifying selection, suggesting important biological roles of the corresponding proteins. Moreover, the functional characterization showed that distinct UGT genes were involved in the biosynthesis of *cis-* and *trans*-isomers of piceid, suggesting that these isomers may play different biological roles in grapevine.

## Introduction

Wood is a complex and highly connected chemical assembly of the sugar polymers cellulose and hemicellulose with the phenolic polymers lignin (Tarasov et al., 2018). This dense network also contains non-structural components named extractives, that include compounds such as terpenoids, phenolic compounds and alkaloids, which are often constitutively produced as defense against biotic and abiotic stresses. For example, a wide array of molecules isolated from extractives in different plant species exhibit interesting antifungal properties (Valette et al., 2017). Grapevine wood is known to be a rich source of polyphenols including stilbenes, reaching up to 114 g/kg of dried stems (Goufo et al., 2020). Stilbenes can be constitutively accumulated, and therefore considered as phytoanticipins, or synthesized *de novo* in response to biotic or abiotic stresses, and are then designated as phytoalexins (VanEtten et al., 1994). Vine canes are known to accumulate significant quantities of stilbenes, up to 10 g/kg dry weight (DW), whereas healthy leaves contain less than 0.5 g/kg of DW (Pavela et al., 2017). However, upon pathogen infection, higher and faster accumulation of stilbenes such as resveratrol, piceid and viniferins in leaves has been associated with increased resistance to *Plasmopara viticola* (downy mildew) and *Erisyphe necator* (powdery mildew) (Schnee et al., 2008; Mijailovic et al., 2022). Similarly, canes from *Vitis sylvestris*, which are tolerant to the dieback-associated fungus *Neofusicoccum parvum*, accumulated more stilbenes oligomers in response to fungal infection than *Vitis vinifera* canes, indicating that wood stilbenes may participate in defenses against wood decay fungi (Labois et al., 2020).

Although wood extractives may play important roles in grapevine defense against pathogens, few studies have explored the genetic determinism of their accumulation. Among them, a metabolic quantitative trait loci (mQTL) analysis in an interspecies cross between *V. riparia* and *V. vinifera* has identified two major loci linked to the accumulation of stilbene dimers and trimers on chromosome 18, albeit without functional analysis of the potential candidate genes (Teh et al., 2019). The aim of this study was therefore to investigate further the genetic basis of grapevine wood extractive composition, using a progeny from a cross between Riesling and Gewurztraminer (Ri x Gw). To this end, we performed an untargeted metabolic profiling of extractives from canes and trunks in the Ri x Gw progeny over 3 years. Genetic analyses led to the identification of a major mQTL linked to the accumulation of resveratrol glucosides in wood tissues. Combined genomic functional and structural analyses highlighted genes encoding uridine diphosphate glycosyltransferases (UGTs) as major determinants of the constitutive accumulation of resveratrol glucosides in grapevine canes and trunks.

## Materials and methods

### Plant material

Genetic analyses were performed on a progeny resulting from the cross of *Vitis vinifera* cv. Riesling clone 49 (Ri) and *Vitis vinifera* cv. Gewurztraminer clone 643 (Gw) (Duchêne et al., 2009). This F1 population, referred to as Ri x Gw, was planted in 2006 in an experimental vineyard at Bergheim (France) (48°21’N, 7°34’E). Between 2020 and 2022, grapevine canes from 103 to 112 offsprings of the Ri x Gw population were collected during winter pruning in December. Canes were kept at 4°C before cutting internodes that were frozen using liquid nitrogen to prevent metabolite oxidation. In April 2022, chips from trunks (about 3 g) of every living offsprings (n=218) were collected, frozen with liquid nitrogen and kept at −20°C.

### Chemicals

LCMS grade methanol and acetonitrile were purchased from Roth Sochiel (France); water was provided by a Millipore water purification system. Substrates for enzyme studies (resveratrol, nerol, quercetin, kaempferol) and analytical standards (piceid, quercetin 3-glucoside and kaempferol 3-glucoside) were purchased from Sigma-Aldrich (L’lsle-d’Abeau, France). *Cis*-resveratrol was obtained following exposing *trans*-resveratrol to daylight. Chloramphenicol (Sigma-Aldrich) was used as internal standard.

### Sample preparation and metabolites extraction

Canes and chips were lyophilized for 24 hours. For each genotype, internodes were roughly grounded with a conventional coffee grinder (Moulinex, France), then a finer grind was carried out using a ball mill (TissueLyser II, Qiagen, Courtaboeuf, France). Five mg ± 0.1 mg of powder was weighted in 2 mL polypropylene microtubes (Eppendorf). A first microwave assisted extraction (Samsung, 450 W, 30s) was performed with 200 µL of LC–MS grade methanol containing 5 mg/L of chloramphenicol as an internal standard, followed by an ultrasound bath for 2 min (Fisherbrand, FB15050). Samples were centrifuged (11 000 g, 10 min, 20 °C). A second similar extraction without microwaves was performed on the pellet, and both methanol extracts were pooled together.

### Untargeted metabolomic analyses

UHPLC-HRMS-based untargeted metabolomic analyses were carried out as described previously (Verdier et al., 2025). Raw data were imported into the Compound Discoverer 3.3.0.550 software (Thermo Scientific, Waltham, MA) and analyzed with an untargeted metabolomics workflow, which included mapping pathways and searching online databases for metabolite annotation. Retention times were aligned to a QC (Quality Control) sample. For compounds detection, a minimal signal-to-noise (S/N) threshold at 1.5 was set as well as a minimum peak intensity of 10 000 and a mass tolerance of 5 ppm. For the gap filling, mass tolerance was set 5 ppm and signal-to-noise threshold at 1.5. The peak annotation and metabolic pathways were performed in comparison to online spectral libraries and online pathways, with a mass tolerance set at 5 ppm. Settings used for the composition prediction were a mass tolerance at 5 ppm. A post-processing filtering was applied, removing ions without formula, Δ-ppm above or below 5 ppm and peak-rating < 4 in QC. The generated ions list with areas was exported as a .csv file. Ion identifiers were generated as MxxxTyyy, where xxx is the *m/z* and yyy the retention time in seconds.

### QTL analysis

Genetic markers consisting of single nucleotide polymorphisms (SNPs) were obtained by a genotyping by sequencing (GBS) as described previously (Arnold *et al*., 2025). QTL detection was performed on both parental and consensus maps of the Ri x Gw population with the R/qtl software (Broman et al., 2003) using the one-dimension scan command scanone. LOD (Logarithm of the odds) significances were ensured with permutation tests (1,000 permutations). The percentage of variance explained by a QTL was assessed with analysis of variance using type III sums of squares using the *fitqtl* function. Confidence intervals were calculated as Bayesian credible intervals using *bayesint* function with a probability of coverage of 0.95.

### Identification of candidate genes

The genome of PN40024.v4 (Velt et al., 2023) was explored to search candidate genes. Physical positions of confidence intervals on PN40024.v4 were given by the physical position associated to SNPs delimiting QTL confidence intervals on genetic maps. Genes between flanking markers were listed with their automatic annotation, and genes without annotation were blasted on NCBI (https://blast.ncbi.nlm.nih.gov/Blast.cgi/). A list of candidate genes was established based on the correspondence between predicted gene functions and metabolites biosynthetic pathways.

### Expression profiles of VvUGTs

Publicly available RNAseq raw data were analyzed using the GREAT tool for comparative transcriptomic analyses (GRapeExpression ATlas; https://great.colmar.inrae.fr/) and heatmaps were generated on Rstudio, using the ggplot2 function.

### Expression of recombinant VvUGTs proteins

VLRSgt sequence was obtained from GenBank (DQ832169) and *VvUGTs* sequences were obtained from the Riesling genome (Zou et al., 2021). Full-length *VvUGT* and *VLRSgt* coding sequences were synthesized and cloned into the pET22b expression vector by the GENEWIZ gene synthesis service (Leipzig, Germany), after codon optimization for expression in *E. coli*. pET22b-UGT plasmids were transformed into BL21 (DE3) competent *E. coli* cells (Sigma-Aldrich, L’lsle-d’Abeau, France). Overnight cultures (2 mL) were inoculated into 50 mL of LB medium containing 1 % (v/v) glycerol and grown at 37°C to an OD_600 nm_ of 1.0, prior to induction with 1mM isopropyl thio-β-D-galactoside (IPTG). Cultures were then grown further for 3h at 37°C. After harvesting the cells by centrifugation (10 min at 6,000*g*), the pellet was resuspended in 3 mL of buffer (50 mM HEPES pH7, 1 mM MgCl_2_, 1 mM β-mercaptoethanol, 1 µg.µL^-1^ bovine serum albumin) and cell lysis was carried out by sonication (10 rounds of 15 sec, with cooling on ice for 30s between rounds). Non-soluble parts were removed by centrifugation (10 min at 13,000 *g*) and *E. coli* extracts were stored on ice before use.

### Enzyme assays

For the determination of kinetic parameters, 50 µL of *E. coli* extracts were incubated with different concentrations of selected substrates (nerol, *trans-*resveratrol, *cis-*resveratrol kaempferol or quercetin) and with 200 µM of UDP-glucose. Incubation buffer was added (50 mM HEPES pH7, 1 mM MgCl_2_, 1 mM β-mercaptoethanol, 1 µg.µL^-1^ bovine serum albumin) to a final volume of 100 µL. Incubations were carried out at 30°C for 30 minutes. Reactions were stopped by heating for 5 minutes at 80°C, followed by addition of 100 µL of LC-MS grade methanol and centrifugation (10 min at 13,000*g*) before LC-MS analysis of reaction products.

K_m_ were calculated from Lineweaver-Burk plots using the Igor Pro 7 software. For V_max_ calculation, the concentrations of recombinant UGT proteins were estimated from SDS-PAGE, compared to a standard range of BSA (0.1 to 4 mg/mL). For substrate specificity studies, *E. coli* extracts were incubated with kaempferol, quercetin and resveratrol at a final concentration of 30 µM in the presence of 200 µM UDP-glucose in incubation buffer in a final volume of 200 µL for 30 min at 30°C. Reaction products analyzed by LC-MS after

### Molecular modelling

UGTs were modeled on the basis of solved crystal structure of Michaelis complexes of VvGT1-UDP-2FGlc Kaempferol (PDB 2C1Z) and PaGT2 UDP-2FGlc resveratrol (PDB 6JEM) (Offen et al., 2006; Maharjan et al., 2020). The UGTs enzyme structures were predicted by Alphafold2 (Jumper et al., 2021). The ligands were manually positioned on the basis of the known Michaelis complexes structures. The modelled complexes were then subjected to molecular simulation. After energy minimization, the complexes were submitted to molecular dynamics during 100 ns with Gromacs (version 2023.1) and the trajectory were analyzed with tools provided by Gromacs (Abraham et al., 2015). In order to control convergence of the simulations, the position of the ligands was analyzed by root-mean-square-displacement analysis (RMSD) from the molecular dynamics trajectory. The binding energy was estimated by free energy calculation using Prodigy-Lig, based on the contacts between the protein and the ligand (Vangone et al., 2019).

### Phylogenetic reconstruction

Relevant UGT sequences from grapevine and *Arabidopsis thaliana* were selected based on the phylogenetic analysis of the grapevine UGT gene family, which includes 228 putative UGTs (Wei et al., 2021). Selected grapevine coding sequences were retrieved from the PN40024.v4 grapevine reference genome sequence (Velt et al., 2023), from the Riesling genome sequence (Zou et al., 2021), and from the *V. rotundifolia* cv. Trayshed genome sequence (Massonnet et al., 2020). Coding sequences from *A. thaliana* were retrieved from TAIR (https://www.arabidopsis.org/). Accession numbers of the coding sequences are shown in **Supplementary Table S7**. Sequence alignment was carried out using MAFFT v7.490 (Katoh and Standley, 2013) with default settings. A maximum likelihood (ML) phylogenetic tree was built using PhyML (Guindon and Gascuel, 2003) accessed through the Seaview graphical interface (http://pbil.univ-lyon1.fr/) (Gouy et al., 2010). The analysis employed the General Time Reversible (GTR) model, incorporating a γ distribution with four rate categories and optimized alpha parameters. To assess node support, 100 bootstrap replicates were performed. For the analyses within the clade of **eight** grapevine UGT proteins, a reduced dataset containing 26 samples was used (**Supplementary Fig. S7)**. For this subset the alignment was carried out using MAFFT v7.490 (Katoh and Standley, 2013) and the phylogenetic tree was built using MrBayes v3.2.7a (Ronquist et al., 2012) with the default settings.

### Selection analyses

dN/dS (or ω) was estimated using both codeml in PAML v4.8a (Yang, 2007) and Fitmodel (Guindon et al., 2004). Codeml analyses were conducted by first implementing branch model and then branch-site models, following the authors’ recommendations. Branch models assume that the ω differs between branches of the tree and was implemented to investigate the selection pressures within the clade of eight UGT proteins. We annotated 8 different foreground branches (i.e. Vitvi16g01943, Vitvi16g01142, Vitvi16g01946, Vitvi16g01150, Vitvi16g01149, Vitvi16g01882, Vitvi16g01143 and Vitvi16g00571), each with an independently estimated ω. Model comparison with the null model (homogeneous ω over all branches) were conducted with Likelihood Ratio Test (LRT) as indicated previously (Álvarez-Carretero et al., 2023). The branch-site model, allowing ω to vary among sites and branches, was then used for the detection of sites under positive selection in specific lineage. Multiple branch-site analyses were conducted with a different branch as foreground. Specifically, we tested for positive selection on sites along the branch ancestral to each of the eight proteins clade and ancestral to proteins that share functional traits. Similarly to branch-models, model comparisons with the null model (i.e. assuming ω=1 for all sites) were conducted using the LRT.

We additionally ran Fitmodel, which accommodates selection shifts across both sites and lineages (Guindon et al., 2004) and was used to explore further possibilities. Following previous studies in grapevine molecular evolution (Parage et al., 2012) we specifically tested the M2a model, which allows for three ω categories (ω1 < ω2 < ω3), and the MX model, which constrains ω categories (ω1 ≤ 0.999; 1.000 ≤ ω2 ≤ 1.100; ω3 ≥ 1.101). We ran both M2a and Mx models allowing switches between selection regimes: +S1, in which the switch between omega categories is identical; and +S2, in which the switch between ω1, ω2 and ω3 is free and can be biased in any direction. LRT statistic was conducted to perform comparison between nested models as indicated previously (Guindon et al., 2004) (i.e., M2a vs Mx; Mx vs MxS1; MxS1 vs MxS2).

### Investigation of amino acid profiles

Based on the results from the selection analysis, the Pelican software v1.0.8 (Duchemin et al., 2023) was used to investigate the amino acid profiles of the proteins of interest. This software aims to detect shifts in amino acid preferences that correlate with a phenotype along a phylogeny, and was previously shown to be suited for investigating episodic changes in selection pressures (Duchemin et al., 2023). Pelican was executed with the ‘*pelican scan discrete’* command using an amino acid alphabet. Differences in the amino acid composition were considered significant when Pelican estimated p-value < 0.05.

## Results

### Untargeted metabolomic analysis of grapevine wood extractives and mQTL analyses

Canes from a segregating population resulting from a cross between *V. vinifera* cv Riesling (Ri) and Gewurztraminer (Gw) were sampled over 3 years (2020, 2021 and 2022) and trunks from the corresponding genotypes were collected in 2022. Metabolic profiling of canes using ultra-high-performance liquid chromatography coupled to high-resolution mass spectrometry (UHPLC-HRMS) yielded between 1459 to 3079 compounds with unique retention time and mass to charge ratio (*m/z*), depending on the year and the type of wood. These compounds were named MxxxTyyy, xxx corresponding to the *m/z* ratio and yyy to the retention time in seconds. mQTL analyses were performed using both the consensus and parental genetic maps. The consensus map consisted of 14428 single-nucleotide polymorphism (SNPs) markers covering 1234.6 cM over 19 linkage groups. Ri and Gw parental maps consisted of 9449 and 9425 SNPs markers respectively, covering 1239 and 1175 cM. QTL analysis using the non-targeted metabolomics data showed that between 339 and 609 compounds (depending on the year and type of wood) were significantly associated to at least one QTL (P < 0.05), with a logarithm of the odds (LOD) score ≥ 3 on the consensus map. Chromosome-level representation of mQTLs (**Fig. 1**) highlighted several hotspots obtained with both the trunks and canes data sets, these hotspots being characterized by a co-localization of a large number mQTLs at a same chromosomal location. The hotspots on chromosome 5 (14-25 cM), 8 (56-59 cM), 10 (10-13 cM), 14 (55-65 cM) and 16 (35-40 cM) were among the most notable. Selection of mQTLs common to all data sets identified a total of 15 common mQTLs, linked to 35 metabolites. The largest number of distinct mQTLs on a single chromosome was 5 for chromosome 18 ( **Supplementary Table S1**). Following detailed analysis of high-resolution mass spectra, putative identification could be obtained for some of these metabolites, which were selected for further investigations. Four metabolites (M345T195, M480T385, M485T385 and M493T380, **Supplementary Table S1**) mapped to a QTL spanning a 540 Kb region on chromosome 5 (3.51 – 4.05 Mb), linked to an allelic variation in Gw. The metabolites M345T195 and M493T380 (chemical formulas C_16_H_26_O_8_ and C_22_H_38_O_12_, respectively) were putatively identified as formic acid adducts of terpenes glycosides. The identity of these ions was also supported by their retention time and their mass spectra, which showed several fragments frequently found in terpene derivatives (*m/z* 59.0138; 71.0138; 89.0243; 101.0246; 113, 131.0487) (Hjelmeland et al., 2015).

**Figure 1.**
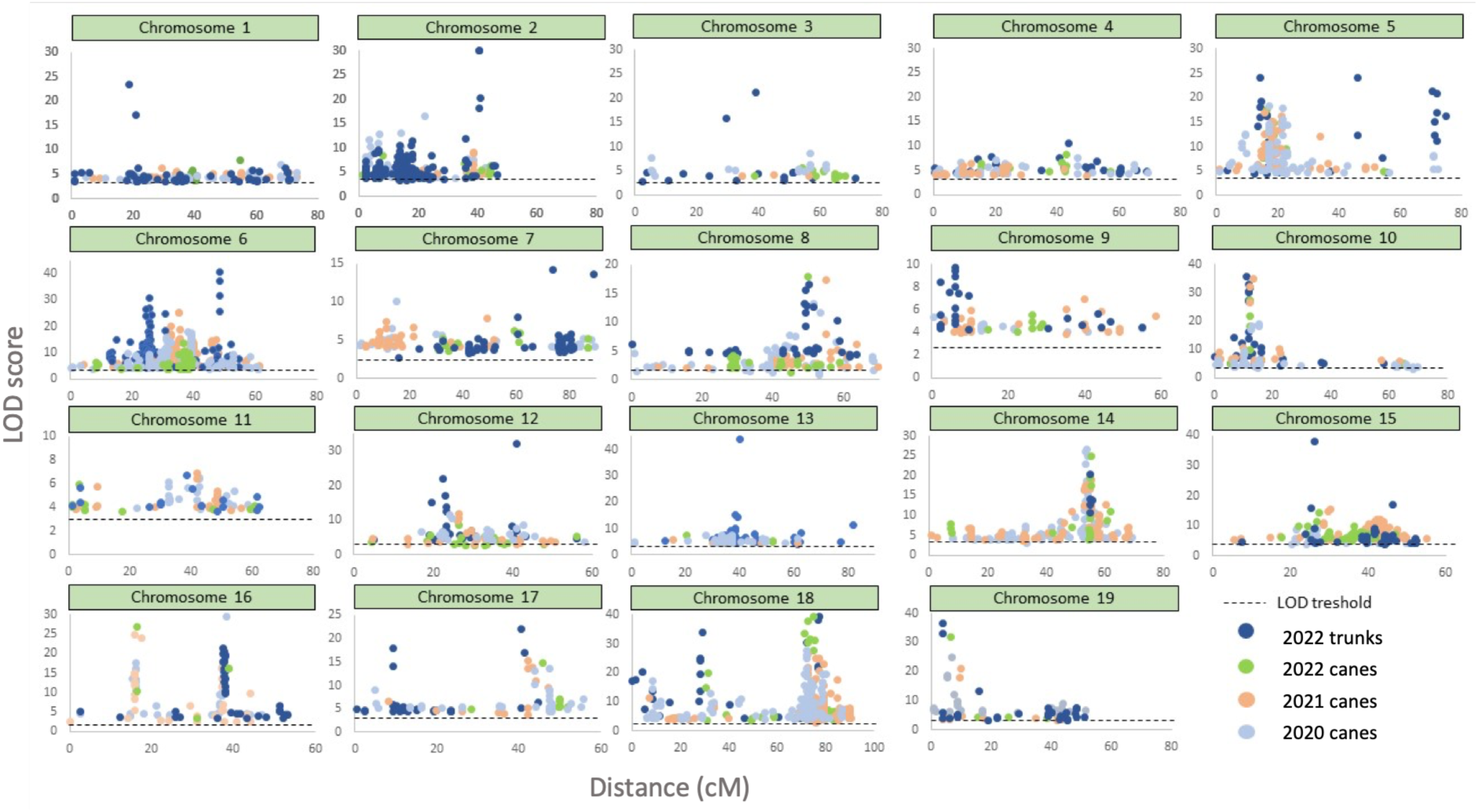
Chromosome-level representation of mQTLs obtained with cane and trunk extracts from Ri x Gw population between 2020 and 2022, based on untargeted metabolomic analyses. Relative quantifications of metabolites from cane and trunk extracts of the Ri x Gw progeny were used to map metabolic QTLs on consensus genetic map, each point corresponding to a significant mQTL (p < 0.05). mQTL positions on chromosomes are shown on the horizontal axis and LOD scores are indicated on the vertical axis.

Three ions, M227T225, M389T225 and M389T308 were associated with the same QTL on chromosome 16 on the Ri map, spanning a region of 1.55 Mb (18.28 – 19.82 Mb). For the M389T225 ion, the LOD score ranged between 13.6 to 17.4, explaining 29.8% to 53% of the phenotypic variance, depending on the data set (**Table 1** and **Fig. 2A**). For the M389T308 ion, the LOD score ranged from 5.4 to 10.7, explaining 19.9-26.5% of phenotypic variance. Finally, the LOD scores associated to the M227T225 ranged from 13.4 to 17.2, explaining 30.4-52.8 of phenotypic variance. Chemical formula associated with M389T308 and M389T225 was C_20_H_22_O_8_, while M227T225 corresponds to C_14_H_12_O_3._ M389T308/M389T225 were identified as resveratrol 3-β-glucoside (piceid), and M227T225 as a resveratrol fragment resulting from in-source fragmentation of piceid. By analyzing retention times and MS² spectra, M389T225 and M389T308 were identified as *trans*-piceid and *cis*-piceid and confirmed using the corresponding commercial standards (**Supplementary Fig. S1**). Segregation of piceid amounts (*trans* + *cis* isomers) in the canes of the Ri x Gw population revealed a continuous variation as illustrated by the example from 2022 (**Fig. 2B**). Comparison of the Ri and Gw parent genotypes, showed that Ri accumulated significantly more piceid than Gw in 2021 and 2022, regardless of the type of wood (p_value_ ≤ 0.05), with this difference not being significant in 2020 (**Fig. 2C**). In addition, comparison of peak areas corresponding to *trans*-piceid and *cis*-piceid indicated that *trans*-piceid accumulated constitutively in greater quantities (approximately five to ten times more) than *cis*-piceid in the wood of the Ri x Gw population (**Supplementary Fig. S2**).

**Figure 2.**
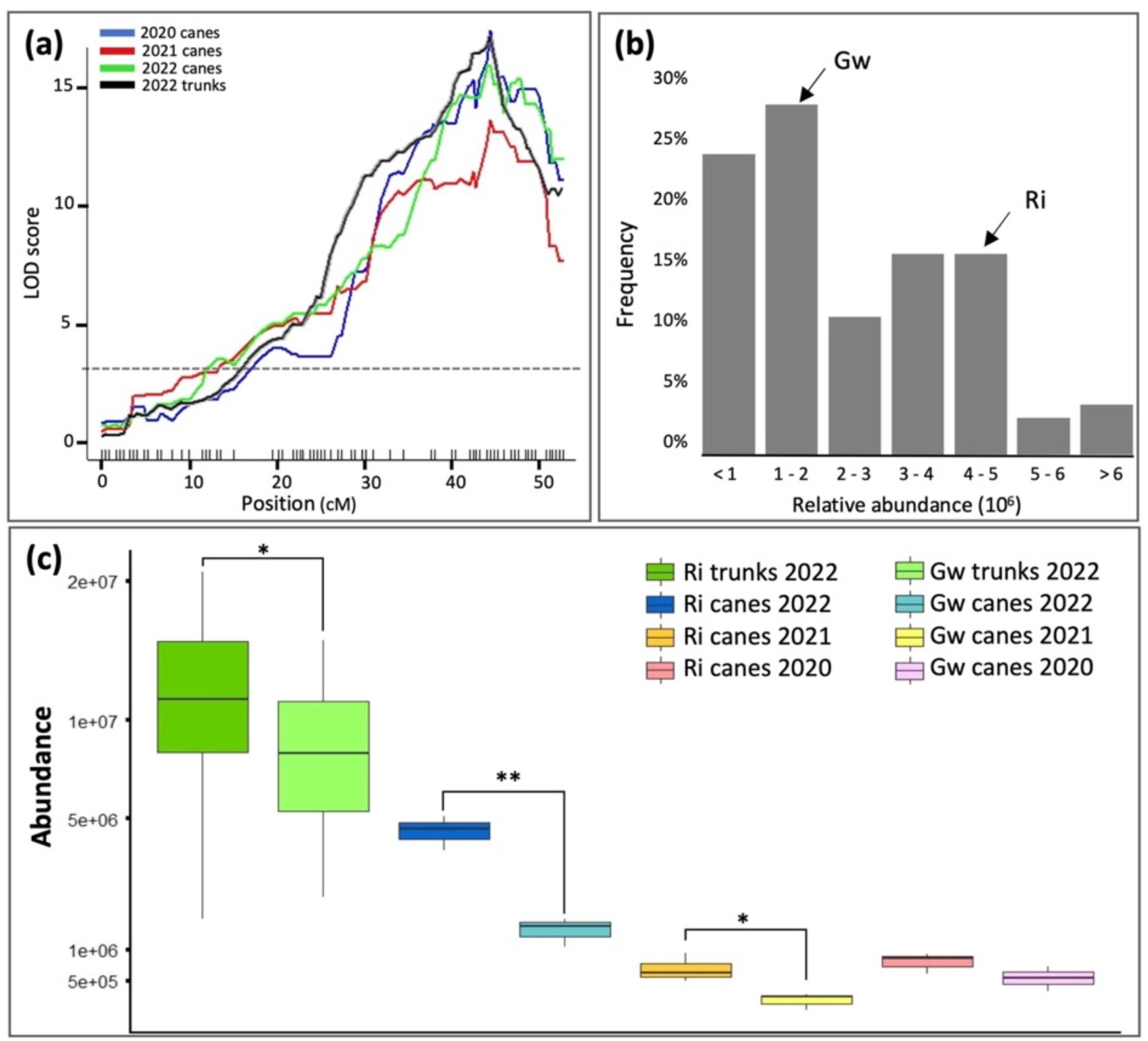
Genetic analysis of piceid accumulation. A) LOD score plots for the metabolite M389T225 (piceid) detected on chromosome 16 on the Ri map (cM) on canes and trunks over 3 years. Dashed gray line shows the threshold of LOD. B) Distribution of relative piceid amounts in 2022 (sum of peak areas of *trans* and *cis* isomers, arbitrary units) in canes of Ri (n = 3), Gw (n = 3) and in the Ri x Gw population (n = 106). C) Relative abundance of piceid (sum of peak areas of *cis* and *trans* isomers) in trunks and canes of Riesling and Gewurztraminer over the 2020-2022 period, * : pvalue < 0.05; ** : pvalue < 0.001.

**Table 1:** Summary table of the mQTLs for piceid accumulation detected in the Ri x Gw population in 2020-2022 for canes and in 2022 for trunks. Chr: chromosome; LOD: Logarithm of the odds; Cm: centimorgans.

| Ion | Formula | Putative identity | Tissues | Year | Map | Chr | LOD score | Peak position (Cm) | Physical position of LOD interval (Mbps) | % variance explained |
| --- | --- | --- | --- | --- | --- | --- | --- | --- | --- | --- |
| M389T308 | $C_{20}H_{22}O_8$ | *cis*-Piceid | Trunks | 2022 | Ri | 16 | 10.7 | 44.47 | 18.13 – 19.83 | 20.2 |
|  |  |  | Canes | 2020 | Ri | 16 | 7.1 | 44.47 | 16.49 – 21.62 | 26.5 |
|  |  |  |  | 2021 | Ri | 16 | 5.4 | 44.47 | 16.49 – 20.04 | 19.9 |
|  |  |  |  | 2022 | Ri | 16 | 6.9 | 44.47 | 17.34 – 20.04 | 26.5 |
| M389T225 | $C_{20}H_{22}O_8$ | *trans*-Piceid | Trunks | 2022 | Ri | 16 | 16.8 | 44.47 | 18.28 – 19.82 | 29.8 |
|  |  |  | Canes | 2020 | Ri | 16 | 17.4 | 44.47 | 18.28 – 19.9 | 53.0 |
|  |  |  |  | 2021 | Ri | 16 | 13.6 | 44.47 | 16.49 – 20.04 | 51.1 |
|  |  |  |  | 2022 | Ri | 16 | 15.9 | 44.47 | 18.13 – 21.56 | 50.8 |
| M227T225 | $C_{14}H_{12}O_3$ | Resveratrol fragment | Trunks | 2022 | Ri | 16 | 17.2 | 44.47 | 18.13 – 19.82 | 30.4 |
|  |  |  | Canes | 2020 | Ri | 16 | 16.3 | 44.47 | 18.28 – 19.9 | 50.7 |
|  |  |  |  | 2021 | Ri | 16 | 13.4 | 44.47 | 18.28 – 20.05 | 42.4 |
|  |  |  |  | 2022 | Ri | 16 | 16.8 | 44.47 | 18.13 – 20.28 | 52.8 |

### Identification of candidate genes potentially involved in piceid biosynthesis

The search for candidate genes potentially linked to the biosynthesis of resveratrol-glucoside was performed within the minimal confidence interval identified on Ri map in trunks and canes, spanning a 1.55 Mbp region on chromosome 16 (18.28-19.82 Mbps). Based on the PN40024.v4 genome sequence (Velt et al., 2023), 141 genes were identified in this region, 98 of which were already annotated. In addition, 25 genes were further annotated by performing Blast searches (https://blast.ncbi.nlm.nih.gov/Blast.cgi/; **Supplementary Table S2**). Analysis of the predicted functions of these 141 genes revealed a small cluster of six putative uridine diphosphate glycosyltransferase (UGT) genes (*Vitvi16g01142, Vitvi16g01143, Vitvi16g01149, Vitvi16g01150, Vitvi16g01943, Vitvi16g01946,* lines highlighted in green in **Supplementary Table S2**). Plant UGTs have been shown to be involved in the glycosylation of a wide array of metabolites (Vogt and Jones, 2000), which made UGT genes good candidates for the synthesis of piceid from resveratrol. Although a few grapevine UGTs have been found to exhibit resveratrol glycosyltransferase activity *in vitro* (Hall and De Luca, 2007; Härtl et al., 2017), none of them was identified based on genetic analyses. Therefore, these UGTs were selected for further investigation of their potential role in piceid biosynthesis *in planta*.

This cluster of six putative UGT genes spanned 90 kb (19.09-19.18 Mbps) on chromosome 16, based on the PN40024_v4 genome (**Supplementary Fig. S3**). The sequences of the corresponding genes in Ri were retrieved from the Riesling genomic sequence (Zou et al., 2021). All Ri UGT genes shared the same structure composed of a unique exon ranging from 1410 to 1443 bp. Comparison of the corresponding protein sequences showed that these UGTs shared between 52% and 95% identity **(Supplementary Fig. S4** and **Supplementary Table S3**). *Vitvi16g01142* and *Vitvi16g01943* were the most similar with 95% identity, while *Vitvi16g01143* was the most distant with less than 60% identity with the other UGTs in the cluster. All these proteins shared a conserved motif of 14 amino acids involved in UDP-sugar recognition (Gachon et al., 2005; Offen et al., 2006). The six genes of this cluster appear phylogenetically related, as they belong to a single clade in the phylogeny of the grapevine UGT family (Wei et al., 2021).

In order to identify more precisely the causal genes among the six candidates, the expression profiles of these UGTs were compared using the GREAT (Grapevine Expression ATlas) transcriptomic analysis tool. Publicly available RNA seq data sets of interest included healthy shoots from Ri, Chasselas blanc and Chasselas Cioutat. RNA seq data sets from Ri and Gw berries were also selected as comparison and summarized on the heatmap **Supplementary Fig. S5**. Although *Vitvi16g01943*, *Vitvi16g01150* and *Vitvi16g01149* exhibited the highest expression levels in shoot tissues, this gene expression analysis did not highlight a specific gene as the main candidate. Therefore, all six genes were selected for functional characterization, together with the previously characterized bi-functional resveratrol/hydroxycinnamic acid glucosyltransferase VLRSgt from *Vitis labrusca*, as a control (Hall & De Luca, 2007).

### Kinetic and substrate preference analysis of the recombinant UGTs

The six UGTs from Ri as well as VLRSgt were expressed as recombinant proteins in *Escherichia coli*. Crude IPTG-induced *E. coli* extracts were used for *in vitro* UGT activity assays, and negative controls were performed with non-induced *E. coli* crude extracts. The catalytic properties of the Ri UGTs and VLRSgt were determined using various candidate substrates (Hall & De Luca, 2007). The monoterpenol nerol was included as a potential substrate, as the Vitvi16g01943 UGT from “Muscat a petits grains blancs” was shown to exhibit terpenol-glucosyltransferase activity (Bönisch et al, 2014). Two of the six UGTs, *Vitvi16g01149* and *Vitvi16g01143* did not display detectable resveratrol glycosyltransferase activity in our conditions, even with longer incubation times. Consequently, these two enzymes have not been included in further assays. Similarly, none of the six UGTs displayed nerol glycosyltransferase activity at any substrate concentration tested (10 - 300 µM) (data not shown).

Incubation of VLRSgt with resveratrol led to the formation of piceid, confirming its resveratrol glycosyltransferase activity. Nevertheless, VLRSgt displayed higher activity and lower Km values with the flavonoids kaempferol and quercetin as substrates (**Supplementary Table S4**). Like VLRSgt, the Vitvi16g01946 enzyme showed glycosyltransferase activity with resveratrol and flavonoids. However, unlike VLRSgt, Vitvi16g01946 showed better kinetic parameters with resveratrol than with flavonoids. Finally, the Vitvi16g01943, Vitvi16g01142 Vitvi16g01150 enzymes appeared as stilbene glycosyltransferases, showing very little or no activity with the tested flavonoid substrates. Vitvi16g01943 and Vitvi16g01142 enzymes both glycosylated resveratrol into piceid, Vitvi16g01943 exhibiting a higher affinity for resveratrol than Vitvi16g01142 as well a higher catalytic efficiency. Vitvi16g01150 enzyme exhibited the highest resveratrol glycosyltransferase activity among the UGTs in the cluster (**Supplementary Table S4**). Its enzymatic parameters were the best of all the enzymes tested, with the lowest K_m_ for resveratrol and the highest turnover rate. The Vitvi16g01150 enzyme did not use the tested flavonoids as substrates.

In order to illustrate substrate preference when different potential substrates are available simultaneously, selected UGT proteins were incubated in the presence of 3 substrates (quercetin, kaempferol and resveratrol, 30 µM each), resveratrol being supplied as a mixture of *trans* and *cis* isomers. In these conditions, different glycosylation patterns were observed (**Fig. 3**). In the presence of both stilbene and flavonoid substrates, both VLRSgt and Vitvi16g01946 and showed a marked preference for the flavonoid substrates (**Fig. 3, B and C**). Glycosylation of kaempferol led to the formation of two isomers of kaempferol-glucoside. Similarly, two or three quercetin-glucoside isomers were produced with these two UGTs. However, when confronted to both stilbene and flavonoid substrates, no piceid was observed with VLRSgt, while Vitvi16g01946 gave rise to little amounts *cis-*piceid. Conversely, Vitvi16g01943 and Vitvi16g01150 did not show flavonoid-UGT activity when incubated with both stilbene and flavonoid substrates. Instead, these UGTs gave rise to *cis*-piceid (Vitvi16g01943, **Fig. 3D**), and *trans*-piceid (Vitvi16g01150, **Fig. 3E**), respectively.

**Figure 3.**
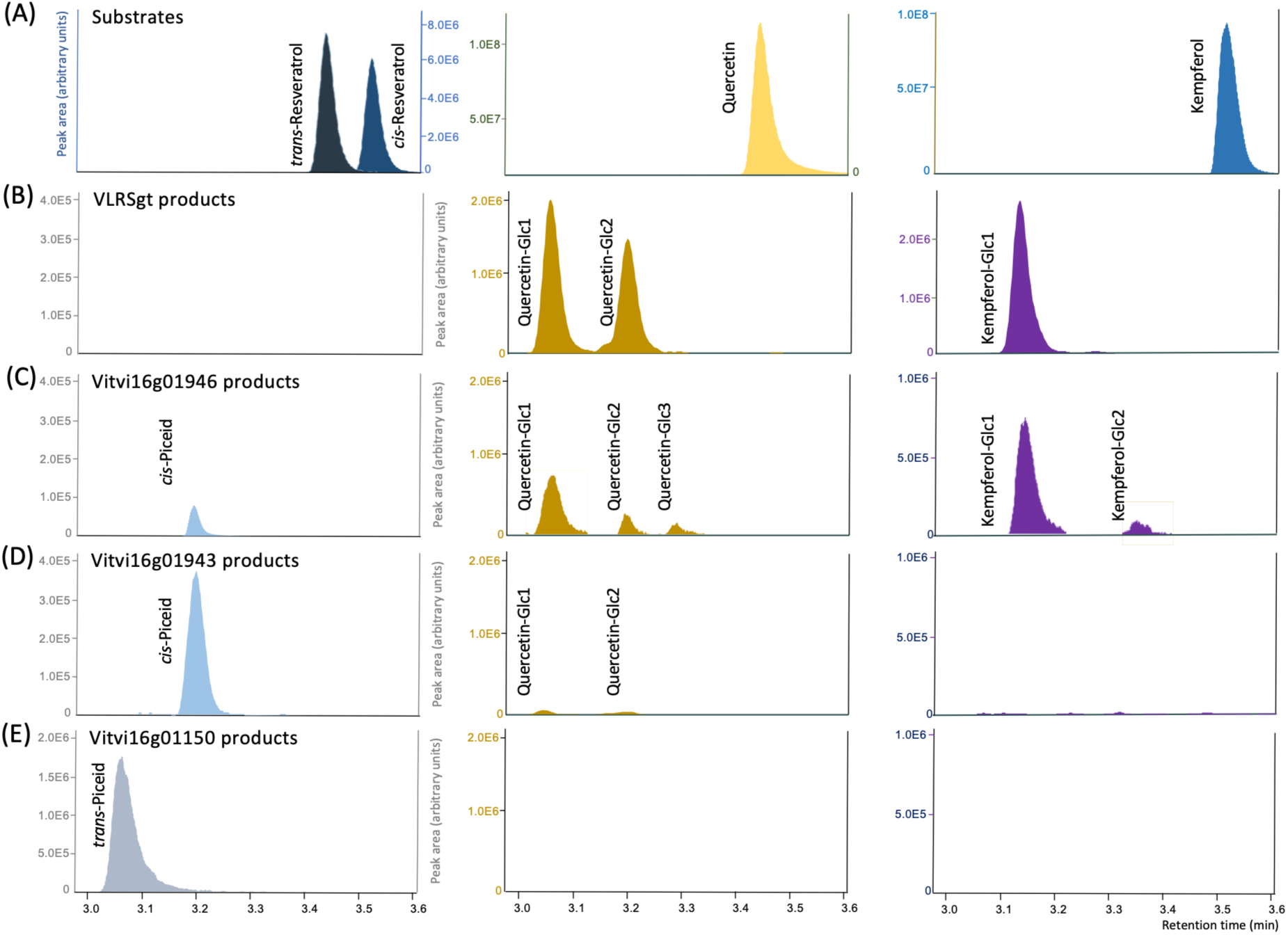
Profiles of products resulting from glucosyltransferase activities with stilbene and flavonoid substrates. LC-MS analysis of reaction products formed by VLRgt (B), Vitvi16g01946 (C), Vitvi16g01943 (D) and Vitvi16g01150 (E), using a mixture of resveratrol, quercetin and kaempferol as substrates (A; 30 µM each), and UDP-glucose as sugar donor (200 µM). Different glucosides of quercetin and kaempferol are numbered Glc1 to Glc3.

### Molecular modeling-based analysis of resveratrol-UGT substrate specificities

Resveratrol was found to be the preferred substrate of both Vitvi16g01943 and Vitvi16g01150 enzymes. However, these UGTs exhibited marked differences in substrate preference for either the *trans* and *cis* isomer of resveratrol, resulting in the specific formation of *cis*-piceid with Vitvi16g01943 and *trans*-piceid with Vitvi16g01150 (**Fig. 3, D and E**). In order to analyze the molecular bases of this regioselectivity and stereospecificity for resveratrol isomers, four Michaelis complexes between these two enzymes, resveratrol (either in *trans* or *cis* configuration) and the sugar donor UDP-glucose (UDPG) were modeled (**Fig. 4**).

**Figure 4.**
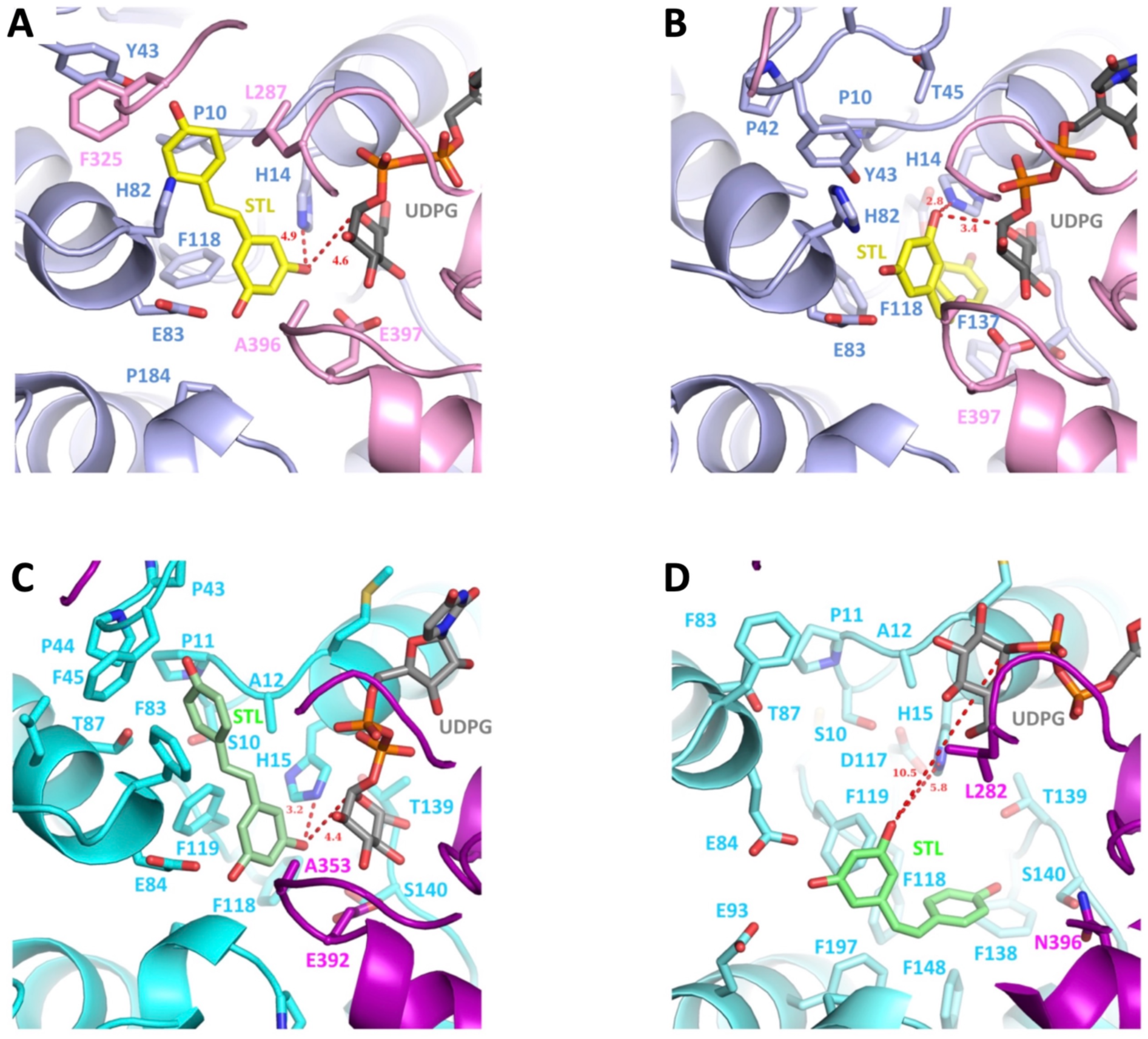
Structural details of the active site of modelled Michaelis complexes between the selected UGT enzymes, resveratrol isomers (STL) and UDP-glucose (UPG). Vitvi16g01943 enzyme with *trans*-resveratrol (A) and *cis*-resveratrol (B). Vitvi16g01150 with *trans*-resveratrol (C) and *cis*-resveratrol (D).

For the complex of Vitvi16g01943 with *cis*-resveratrol, relatively short distances were calculated between resveratrol O3 and Nδ (or Nε depending of the protonation state of imidazole ring) of the catalytic histidine H14, and between resveratrol O3 and C1 of UDPG. These distances were longer in the complex with *trans*-resveratrol (**Fig. 4, A and B**). For Vitvi16g01150 the situation was reversed (**Fig. 4, C and D**), complexes with *trans*-resveratrol showing shorter distance between resveratrol O3, UDPG C1 and the catalytic H14. Furthermore, molecular dynamics simulations predicted lower Kd values for Vitvi16g01943 with *cis*-resveratrol than with *trans*-resveratrol, and lower Kd values for Vitvi16g01150 with *trans*-resveratrol than with *cis*-resveratrol (**Supplementary Table S5**).

### Phylogenetic and molecular evolutionary analysis of the UGT gene cluster on Chr16

Analyses of enzyme activities and substrate preferences within the small UGT cluster on chromosome 16 revealed marked functional diversity. Vitvi16g01946 preferentially glycosylated flavonoids, whereas Vitvi16g01943, Vitvi16g01142 and Vitvi16g01150 were more active on stilbene substrates, with contrasting preferences for trans- and cis-isomers. Nevertheless, these genes belong to the same subfamily of eight phylogenetically related UGTs (Wei et al., 2021), raising the question of whether specific selection patterns accompanied the evolution of substrate preferences.

The maximum likelihood phylogenetic tree **(Fig. 5**) closely matched the topology reported by Wei et al. (2021), with strong bootstrap support (>90%) for most branches. The six UGTs associated to the piceid mQTL clustered together with Vitvi16g01882 and Vitvi16g00571, which did not belong to mQTL confidence interval. A sequence alignment **(Supplementary Fig. S6)** and a phylogenetic tree restricted to this clade **(Supplementary Fig. S7)** were generated for molecular evolution analyses. Selection pressures were investigated using codeml (Yang, 2007) and Fitmodel (Guindon et al., 2004). Branch model analyses with codeml supported heterogeneous selection regimes across the phylogenetic tree indicating specific estimation of ω for the eight genes **(Supplementary Fig. S8)**. Yet, all estimated dN/dS ratios remained below 1, ranging from ω=0.1909 for Vitvi16g01946 to ω=0.6888 for Vitvi16g00571. Branch-site analyses detected significant positive selection only in ancestral branches leading to the (Vitvi16g01943 + Vitvi16g01142) clade and to Vitvi16g01150 (p-values = 0.023 and 0.039, respectively), affecting a very small proportion of sites (i.e. 0.006 and 0.002, respectively).

**Figure 5.**
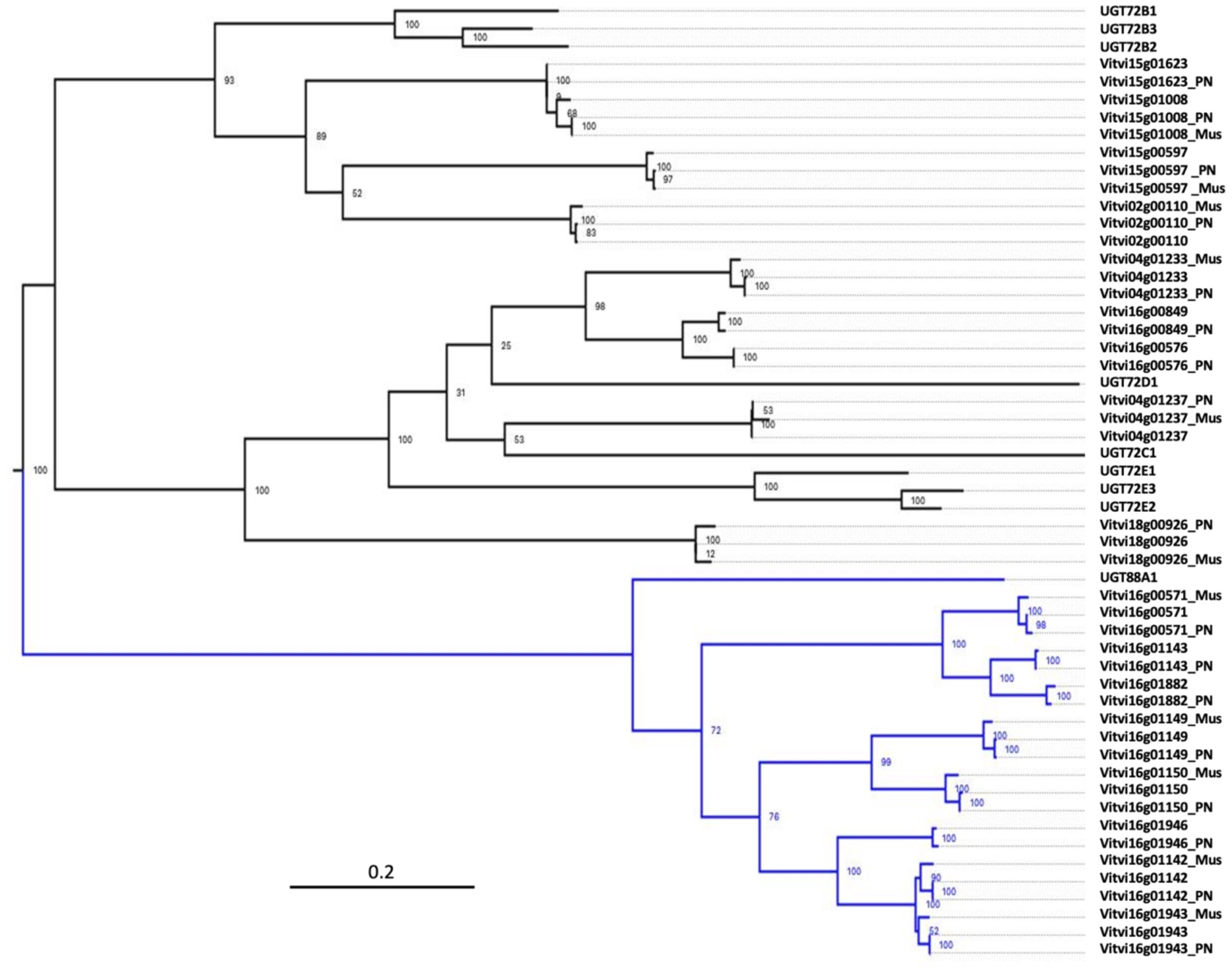
ML phylogenetic tree of UGT coding sequences. The tree was built using 100 bootstrap replicates to assess the robustness of the inferred relationships. Bootstrap support values are displayed on the tree. The clade highlighted in blue was used in downstream analyses and includes the Chr16 clustered UGT genes. Branch lengths are proportional to the number of substitutions per site, as indicated by the scale bar.

Fitmodel analyses identified the Mx+S2 model as the best fit (ω1=0.001, ω2=1.000, ω3=2.481; p=0.002), confirming pervasive purifying selection, with 370 to 403 sites evolving under constraint depending on the gene. No positively selected sites were detected, although relaxed selection was identified at site 65 in the ancestral branch of the (Vitvi16g01943 + Vitvi16g01142) clade **(Supplementary Fig. S8)**.

To investigate potential links between sequence evolution and functional divergence, Pelican analyses (Duchemin et al., 2023) were conducted using either exploratory comparisons (by comparing each protein to all the others to highlight possible individual specificities) or predefined functional categories based on substrate specificity and enzymatic activity. Twenty-eight sites showed significant differences in selection pressures associated with functional characteristics (p<0.05) **(Supplementary Table S6)**. Twenty-four were located in peripheral protein regions, including four sites (32, 60, 76 and 166) associated with resveratrol glycosyltransferase activity **(Supplementary Fig. S9)**. The four remaining sites (16, 17, 22 and 125) were located near the active site **(Fig. 6A**). Sites 16 and 22 were associated respectively with Vitvi16g01150 and with proteins displaying resveratrol glycosyltransferase activity, and both showed relaxed selection linked to proline loss (Fitmodel results). In contrast, site 17 evolved under purifying selection within resveratrol glycosyltransferases proteins, with conservation of a methionine residue. Site 125 was specifically associated with enzymes preferring cis-resveratrol substrates (Vitvi16g01943 and Vitvi16g01142), where relaxed selection is associated with the loss of a conserved phenylalanine (Phe) residue. The significant steric difference between Phe, which is associated with trans-resveratrol preference, compared to smaller residues (e.g. Cys, Ser, Gly) associated with cis-resveratrol preference, suggested a role for this site in substrate specificity.

**Figure 6.**
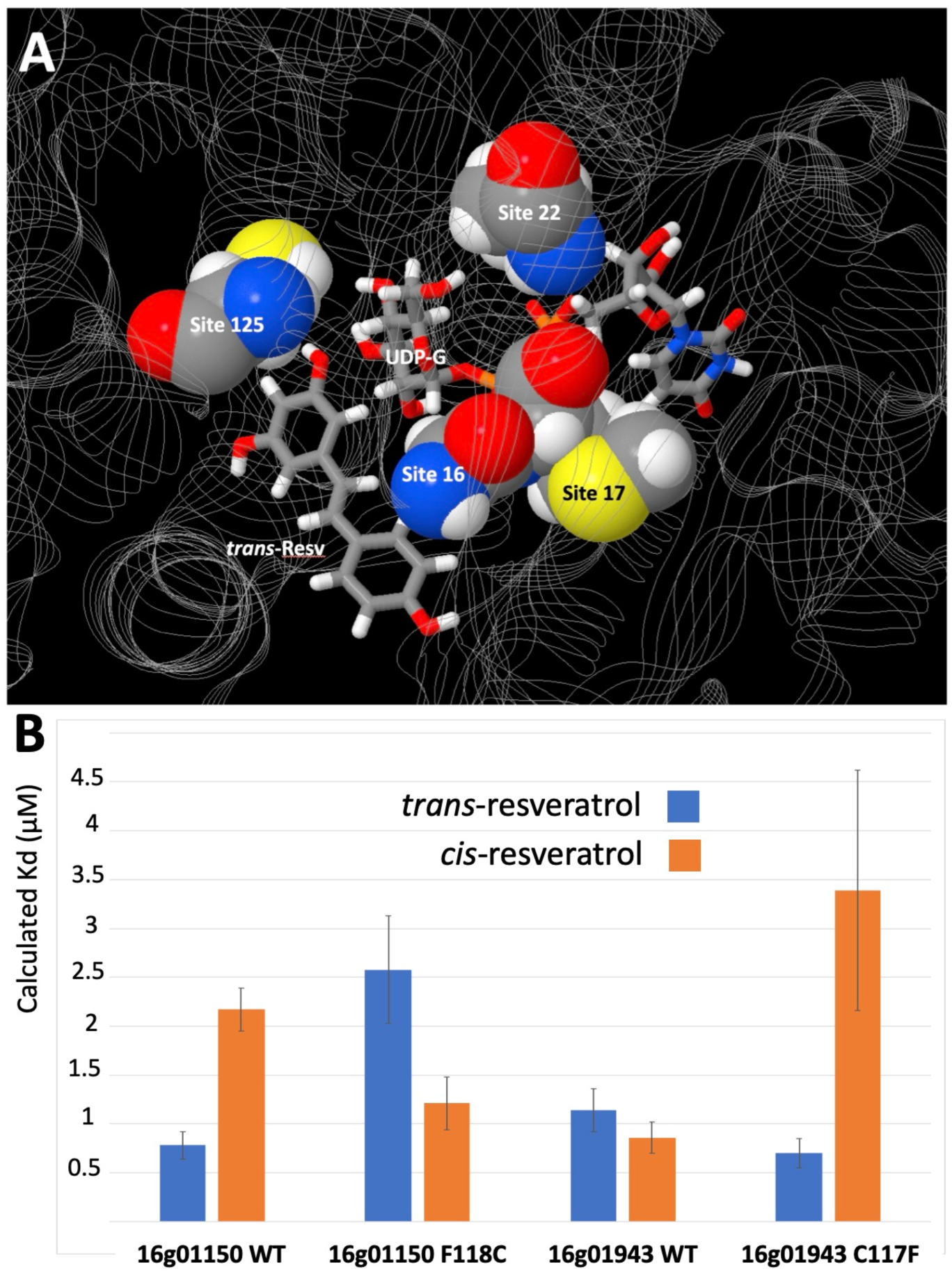
A) Positions of amino acids highlighted by the Pelican analysis and located near the active site of the UGT enzymes. Sites are illustrated in the Vitvi16g01943 structural model with *trans*-resveratrol (*trans*-Resv) and UDP-glucose (UDP-G) substrates. B) Calculated impact of the amino acid at site 125 on substrate preference for *trans*- or *cis*-resveratrol of the Vitvi16g01150 and Vitvi16g01943 enzymes. Kd (μM) were calculated from Molecular Dynamics trajectories (100ns).

Molecular modelling has been shown to accurately predict the impact of mutations on the activity of UGT enzymes (Gerloff et al., 2023); it was therefore used to evaluate the impact of different amino acids at position 125 on substrate affinity. As suggested above, exchanging Phe and Cys at this position in Vitvi16g01150 and Vitvi16g01943 WT models altered their respective calculated affinities for *trans*- and *cis*-resveratrol, reversing optimal substrate affinities in both enzymes **(Fig. 6B**).

Finally, the evolution of distinct UGT genes for *cis-* and *trans*-piceid biosynthesis in grapevine suggest that these isomers may have different biological roles, which may be reflected by different regulation patterns of the corresponding genes. Expression analysis of the *Vitivi16g01150* and *Vitivi16g01943* genes in 2109 publicly available RNA-seq samples showed indeed different expression patterns. Both genes were expressed in most tested organs. However, expression of *Vitivi16g01150* was higher than that of *Vitivi16g01943* in roots, whereas expression of *Vitivi16g01943* was highly dominant in ripening berries (**Supplementary Fig. S10**).

## Discussion

### Genetic analysis wood extractive composition highlights hotspots of mQTLs on the grapevine genome

Despite the ecological and economic importance of wood chemical components, few studies have explored the genetic bases of wood extractive composition (Stackpole et al., 2011; Ganthaler et al., 2017). In this work, we have used a non-targeted metabolomic approach to investigate the genetic bases of grapevine wood extractive composition. Metabolomic analyses were performed on canes of the Ri x Gw progeny for three consecutive years. Trunk extractives were analyzed only in 2022, as the Ri x Gw progeny was uprooted and phenotyped for esca disease damages (Arnold et al., 2025). The genetic analysis wood extractive composition in both canes and trunks has highlighted mQTLs distributed on all grapevine chromosomes, some of them with high LOD scores (**Fig. 1**). Several genomic regions impacted the accumulation of large numbers of metabolites and were thus identified as hotspots of mQTLs. For example, the mQTL hotspot identified on chromosome 5 impacted the accumulation of 20 to 56 metabolites, depending on the year and the tissue considered. Many of these metabolites have been identified as terpene derivatives, indicating that this QTL corresponded to the previously characterized 1-deoxy-D-xylose-5-phosphate synthase (DXS) gene (Battilana et al., 2009; Duchêne et al., 2009). Nevertheless, these results indicated that the Gw DXS allele linked to this QTL not only impacted the accumulation of terpenoids in grape berries as shown previously in the RixGw population (Duchêne et al., 2009), but had also major effects on terpenoid profiles in woody tissues. Beside the mQTL hotspots, some mQTLs were linked to very few metabolites, suggesting that the corresponding genomic regions were involved in the specific control of a small number of compounds. Analysis of high-resolution mass spectra identified two metabolites underlying a stable QTL on chromosome 16 as *trans-*piceid and *cis*-piceid. No other glycosylated compounds mapped this genomic region, suggesting that the genes underlying this mQTL were specifically involved in resveratrol metabolism. Although QTLs linked to stilbene derivatives have been described previously on chromosomes 11, 14 and 18 in a F2 population derived from a *V. riparia* hybrid, no functional characterization of related candidate genes has been performed (Teh et al, 2019). Therefore, the mQTL corresponding to piceid was selected for further investigation.

### A cluster of putative UDP glycosyltransferases co-localizes with a mQTL of resveratrol glucoside

In *V. vinifera*, chromosome 16 harbors a large cluster of 42 stilbene synthase (*STS*) genes (Parage et al., 2012; Vannozzi et al., 2012). Although STS enzymes are involved in the biosynthesis of resveratrol, which is a direct precursor of piceid, the genomic region containing the *STS* gene cluster did not colocalize with the piceid mQTL. In addition, two QTL linked to *cis-*piceid and astringin (piceatannol-3-β-glucoside) were reported on chromosome 16, with candidate genes including *MYB* transcription factors and *STS* (Vezzulli et al., 2019). However, the genomic regions underlying these two QTLs did not colocalize with the piceid mQTL identified in this work.

Interestingly, the piceid mQTL co-localized with a cluster of six genes encoding putative UDP glycosyltransferases, which were selected as candidate genes based on their predicted function. Indeed, plant UGTs catalyze sugar transfer from UDP-activated sugar donors to a wide variety of substrates, ranging from endogenous metabolites to xenobiotics, affecting their biological activity, solubility, stability and transport capacities. Grape UGTs constitute a large gene family, with at least 228 members in the grapevine genome (Ono et al., 2010; Wei et al., 2021). To date, few UGTs exhibiting resveratrol glycosyltransferase activity *in vitro* have been identified in *Vitis* species. The *VLRSgt* gene was identified following the purification of the corresponding protein from the mesocarp of *Vitis labrusca* berries. The VLRSgt enzyme was characterized as a bi-functional resveratrol/hydroxycinnamic acid glucosyltransferase, albeit showing a very significant activity with the flavonoid kaempferol (Hall and De Luca, 2007). Based on the PN40024 reference genome, the orthologue of the *VLRSgt* gene in *V. vinifera* is likely to be the *Vitvi03g00557* gene, with 99.6% identity at the protein level. More recently, *UGT72B27* (*Vitvi15g01623)* has been identified as resveratrol/guaiacol glucosyltransferase involved in the glycosylation of smoke-derived volatiles (Härtl et al., 2017). In the present analysis of wood extractives, none of the genomic regions comprising the *Vitvi03g00557* (orthologue of *VLRSgt)* or the *Vitvi15g01623 (UGT72B27*) genes were associated to a mQTL of resveratrol glucoside. Nevertheless, the VLRSgt enzyme (GenBank DQ832169, Hall & De Luca 2007) was selected as a positive control for resveratrol glycosyltransferase activity *in vitro*.

### *Vitvi16g01150* is a key gene in the constitutive accumulation of piceid in grapevine canes and trunks

Gene expression analyses showed that all genes in the cluster were expressed in woody tissues, without highlighting a particular UGT as the best candidate for piceid biosynthesis (**Supplementary Fig. S5**). Therefore, all genes were expressed in *E. coli* for functional characterization of the corresponding proteins (**Supplementary Table S4**, **Fig. 3**). The Vitvi16g01943 enzyme was previously described as a monoterpenol-glycosyltransferase with affinity for nerol, geraniol and citronellol (Bönisch et al., 2014). However, Vitvi16g01943 from Ri did not show any glucosyltransferase activity with nerol in our experiments, suggesting that the enzyme corresponding to the Ri allele tested here is not active with monoterpenols. In contrast, the Vitvi16g01943 enzyme used resveratrol as a substrate, producing a majority of *cis*-piceid. Kinetic parameters were relatively close to those of UGT72B27 with *trans*-resveratrol as a substrate (K_m_=36 µM, K_cat_=0.61 s^-1^) (Härtl et al., 2017). Although Vitvi16g01142 was very similar to Vitvi16g01943 (95% sequence identity at the protein level), it displayed a lower affinity and kinetic efficiency with resveratrol, consistent with the fact that similar UGTs can have different substrate preferences (Yin et al., 2016; Maharjan et al., 2020).

Finally, only the Vitvi16g01150 enzyme produced almost exclusively *trans-*piceid from resveratrol (**Fig. 3**), in accordance with the pattern of piceid isomers found in grapevine canes and trunks. With resveratrol as a substrate, the Vitvi16g01150 enzyme exhibited better kinetic parameters than the other UGTs characterized in this work. The Vitvi16g01150 enzyme did not show significant activity with quercetin and kaempferol, suggesting a marked specificity for stilbene substrates. The relatively high expression levels of the *Vitvi16g01150* gene in grapevine canes (**Supplementary Fig. S5**) and the high efficiency of the corresponding enzyme with *trans*-resveratrol provide strong evidence in favor of the major role of *Vitvi16g01150* in constitutive piceid biosynthesis grapevine canes and trunk, considering that *trans*-piceid is the major isomer accumulated in these tissues (**Supplementary Fig. S2**). Allelic variations between Ri and Gw are probably responsible for differential piceid accumulation in the woody parts of these two varieties.

### Distinct UGTs are involved in the biosynthesis of *cis* and *trans* isomers of piceid

Grapevine has been shown to accumulate both *cis* and *trans* isomers of piceid, albeit in different amounts depending on tissues and circumstances. For example, various amounts of both *cis-* and *trans*-piceid have been detected in berry skin and wines from different grape varieties, *cis-*piceid often being the major form of stilbene glycoside (Romero-Perez et al., 2001). Similarly, both *cis-* and *trans*-piceid accumulate in grapevine leaves in response to *Plasmopara viticola* infection (Chitarrini et al., 2017). Interestingly, *cis*-piceid was found to be the main stilbene accumulated *V. amurensis* leaves in response to UV-C light (Kiselev et al., 2017), as well as in *V. vinifera* leaves submitted to LED light (Ahn et al., 2015). In the present work, canes of Ri and Gw contained both *cis-* and *trans*-piceid, with a large predominance of *trans*-piceid. On the other hand, expression of a stilbene synthase gene from sorghum in transgenic *Arabidopsis thaliana* resulted in the exclusive accumulation of *cis*-piceid, indicating a marked preference of Arabidopsis UGTs for cis-resveratrol (Yu et al., 2006).

Differential accumulation of *cis* and *trans* isomers of piceid in various tissues of several plant species suggests that different UGTs may be responsible for the biosynthesis of *cis* and *trans*-piceid isomers *in planta*. This hypothesis was supported further by the different substrate and product specificities of some of the UGTs characterized in this work. In order to analyze the molecular bases of the regioselectivity and stereospecificity for resveratrol isomers of the UGTs Vitvi16g01943 and Vitvi16g01150, molecular models of these enzymes were confronted to *cis* and *trans*-resveratrol substrates. Based on these molecular models, calculated distances between the catalytic histidine H14, resveratrol O3 and C1 of UDPG varied significantly depending on the UGT and the resveratrol isomer (**Fig. 4**). Shorter distances, compatible with efficient catalysis, were obtained with *cis*-resveratrol for Vitvi16g01943 and with *trans*-resveratrol for Vitvi16g01150, in agreement with the *in vitro* substrate and product specificity of these enzymes. In Vitvi16g01150, binding of resveratrol in *trans* conformation was favored. The B-ring was stabilized by F45, F83 and F119, P11 and A12. F118 interacted with the A-ring and *trans*-resveratrol was therefore stabilized in optimal position for efficient catalysis, with short distances between atoms involved in catalysis. Conversely, the distances calculated with *cis*-resveratrol were very long. Altogether, the contrast between *trans* and *cis*-resveratrol was the highest for Vitvi16g01150, consistent with the marked preference of this enzyme for *trans*-resveratrol *in vitro*. Sequence analysis showed that Vitvi16g01150 has a Thr residue in helix α3 (T86 and T87 respectively), while Vitvi16g01943 has an Ala. Vitvi16g01150 has also a Ser at position 88, which is replaced by a Phe is Vitvi16g01943. This more polar environment in Vitvi16g01150 may favor the binding of the *trans* isomer of resveratrol in a position compatible with catalysis. Altogether, these different substrate binding characteristics may be explained by different structures of the ligand binding pockets pocket, defined by a flexible loop (loop between β3 and α3). This flexible loop, which is divergent among UGTs, has been predicted to modulate substrate specificity (Wang, 2009).

### Molecular evolution analyses highlight a specific amino acid potentially associated to the evolution of distinct UGTs for *cis* and *trans*-piceid biosynthesis

The topology of the obtained phylogenetic tree, together with the physical distribution of the eight related UGT genes, suggest that these paralogs originated through tandem duplication events, followed by subsequent translocations of Vitvi16g01882 and Vitvi16g00571 to other genomic locations (1.4 Mb and 9.1Mb from the cluster of 6 UGTs, respectively). Gene duplication is assumed to be a major driving force in the evolution of the extraordinary diversity of plant secondary metabolism (Pichersky and Gang, 2000). Like in other gene families involved in plant metabolism, gene duplications followed by the evolution of regions involved in substrate specificity have contributed to the creation of the current functional diversity of UGTs. For example, natural and artificial selection is likely to have resulted in evolution and functional divergence of the UGT85 group in tea (*Camellia sinensis*), which shows substrate preference for monoterpenes, diterpenes and apocarotenoids (Wang et al., 2025). Our dN/dS analyses indicate that the UGT genes considered in this work are overall evolving under purifying selection. The few sites inferred to be evolving under positive or relaxed selection were detected in ancestral branches of *cis*- and *trans*-stilbene-specific lineages (sites 161, 462 and 65, **Supplementary Fig. S8**).

A total of 28 sites showing significant differences in selection pressures depending on functional characteristics (p-value < 0.05) were identified using Pelican **(Supplementary Table S6)**. Of these, 24 were located in peripheral protein regions, while 4 were located near the active site. Among the 24 peripheral sites, four (positions 32, 60, 76, and 166) showed distinct evolutionary patterns in proteins with resveratrol glycosyltransferase activity and may therefore be associated with the emergence of this function. Interestingly, among the four sites located near the active site, two sites (positions 17, methionine gain, and 22, proline loss) were also identified in enzymes displaying resveratrol glycosyltransferase activity. Together, these results suggest that the establishment of this function may have involved coordinated changes affecting both peripheral regions and the active site.

The remaining active-site-proximal sites provided further insight into substrate specificity. In particular, site 16 identified in protein Vitvi16g01150, which shows a preference for *trans*-resveratrol, may contribute to substrate affinity through the loss of a proline residue. Site 125 is also noteworthy: the Fitmodel analysis detected relaxed selection associated with decrease in conservation of a phenylalanine residue at this position, specifically in enzymes with a preference for *cis*-resveratrol (e.g., Vitvi16g01943 and Vitvi16g01142). Molecular modelling further supported its functional relevance. Indeed, exchanging the residues from Phe to Cys at this position (corresponding to site 125, i.e., F118 in Vitvi16g01150 WT and C117 in Vitvi16g01943 WT) affected the predicted affinities for *trans*- and *cis*-resveratrol substrates in both enzymes, leading to a reversal of substrate preference (Fig. 6B). The marked difference in steric hindrance between Phe, associated with *trans*-resveratrol preference, and smaller amino acids (Cys, Ser, Gly), associated with *cis*-resveratrol preference, as well as the calculated impact of these amino acids on enzyme affinity, suggest that this site may have played a key role in the evolution of distinct UGTs for *cis* and *trans*-piceid biosynthesis. In plants, a small number of key amino acids can strongly influence the substrate preference and stereospecificity of UGT enzymes (Del Giudice et al., 2022). Such changes likely facilitated the functional divergence of UGT gene families and contributed to the diversification of specialized metabolism in plants (Han et al., 2024; Xu et al., 2026). Similarly, the diversification of substrate preferences within the small UGT cluster on chromosome 16 may have contributed to the structural diversification of stilbenoids in grapevine.

Although the role of wood extractives in defense against fungal pathogens has been demonstrated (Labois et al., 2020), relatively few studies have aimed to decipher the genetic determinism of their biosynthesis. Following a genetic analysis of grapevine wood composition, this study represents the first report on the identification and functional characterization of genes involved stilbene biosynthesis using a quantitative genetic approach. Based on the characteristics of the corresponding enzyme, *Vitvi16g01150* appears to be the primary gene involved in the constitutive accumulation of *trans*-piceid, the most abundant stilbene glycoside in grapevine canes and trunks. However, the functional characterization of the neighboring *Vitvi16g01943* gene revealed a marked preference of the corresponding enzyme for the *cis*-resveratrol substrate, suggesting that this gene is specifically involved in the biosynthesis of *cis*-piceid. Future work will be aimed at deciphering the biological significance of the emergence of specific glycosyltransferases involved in the biosynthesis of *cis*- and *trans*-piceid, respectively.

## Acknowledgments

We are grateful to our colleagues of the Unité Expérimentale Agronomique et Viticole (UEAV) at INRAE-Colmar (France) for maintaining the vineyard plots. We thank Fabrice Neiers and Mathieu Schwartz (CSGA, Dijon, France) for valuable advice about optimization of the gene synthesis and cloning procedure into the pET22b vector. This work was supported in part by the “Fonds régional de coopération pour la recherche” of the Grand Est region (France) through the VitEst project. HC was supported by a fellowship from the French Ministry of Research and Higher Education. The funders had no role in study design, data collection and analysis or preparation of the manuscript.

## Author contributions

HC, RT, AMG and PH designed the research, interpreted the data and wrote the manuscript, with input from all authors. HC, GR and AMG performed wood sampling and metabolomic analyses; HC, GB, VD, GA, KA and ED participated in genetic analyses; HC, ML, AV and CR performed genomic analyses; SR conducted protein modeling analyses; HC, JFC and PH performed enzyme kinetics experiments; CB, ZP and RT conducted evolutionary analyses; PH supervised the study and organized the acquisition of fundings.

