## Supplemental Figs and tables for "Genetic analysis of grapevine wood composition identifies glycosyltransferases involved in the biosynthesis of stilbene glycosides"

### Supplementary data

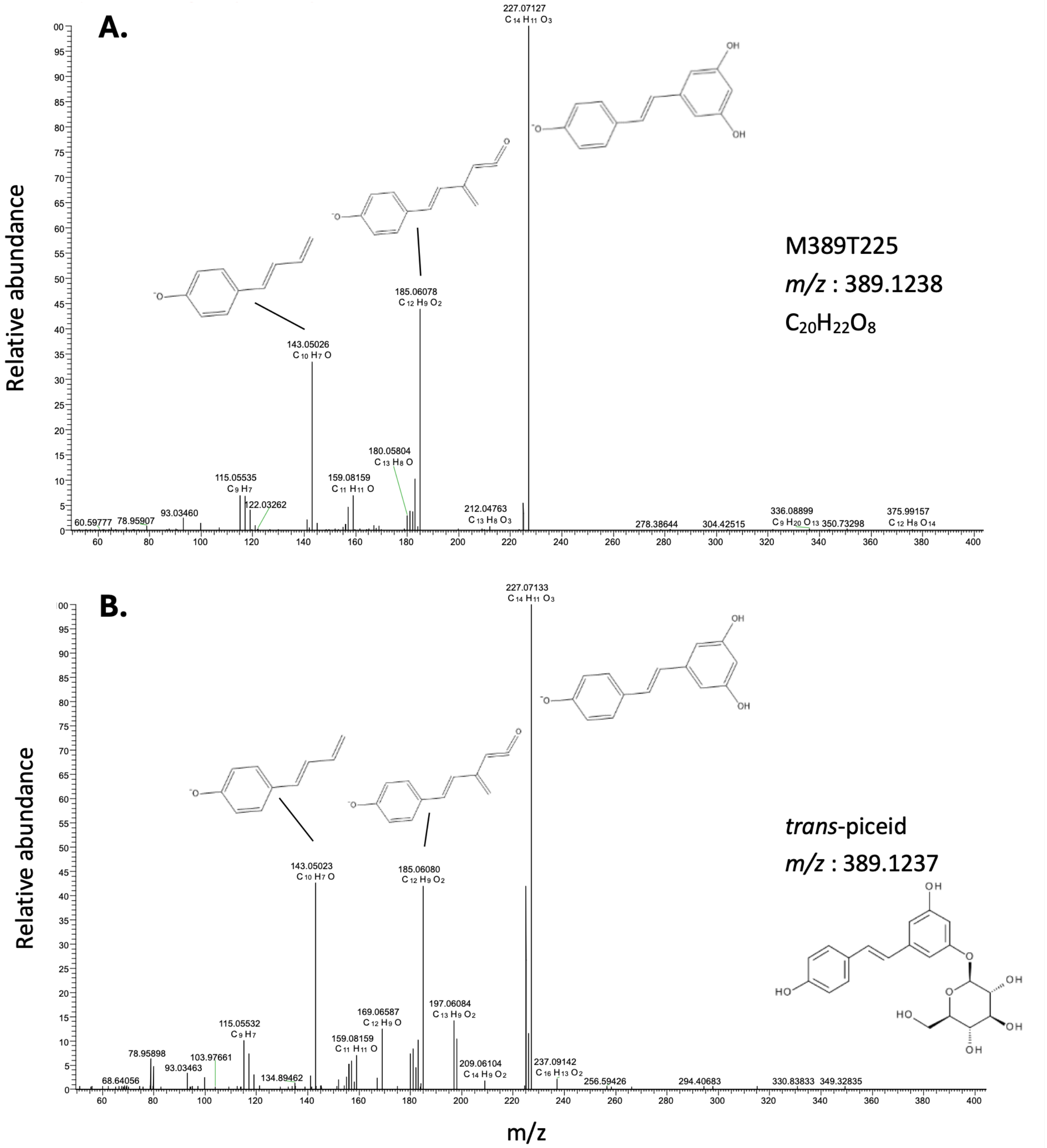

**Figure S1.** MS/MS spectra of M389T225 (A) and *trans*-piceid standard (B) in negative ionization mode.

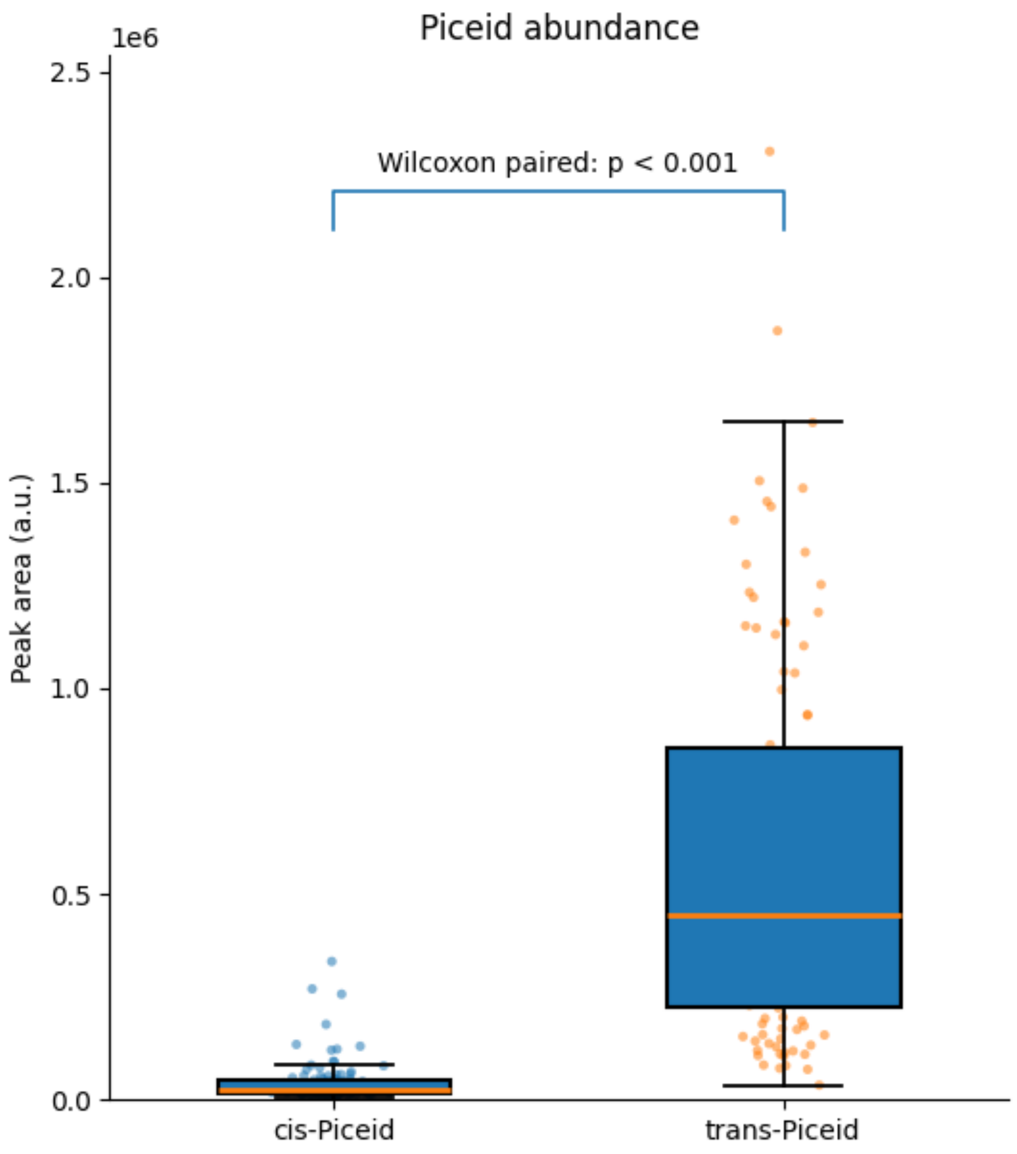

**Figure S2**. Relative amounts of *cis*-and *trans*-piceid in the canes of the Ri x Gw population (2021)

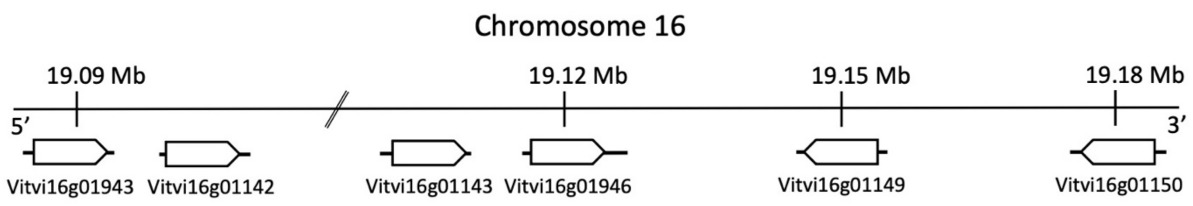

**Figure S3**. Genomic structure of the UGT gene cluster on chromosome 16, based on the PN40024_v4 genome sequence. Boxes indicate the coding sequences of the indicated genes.

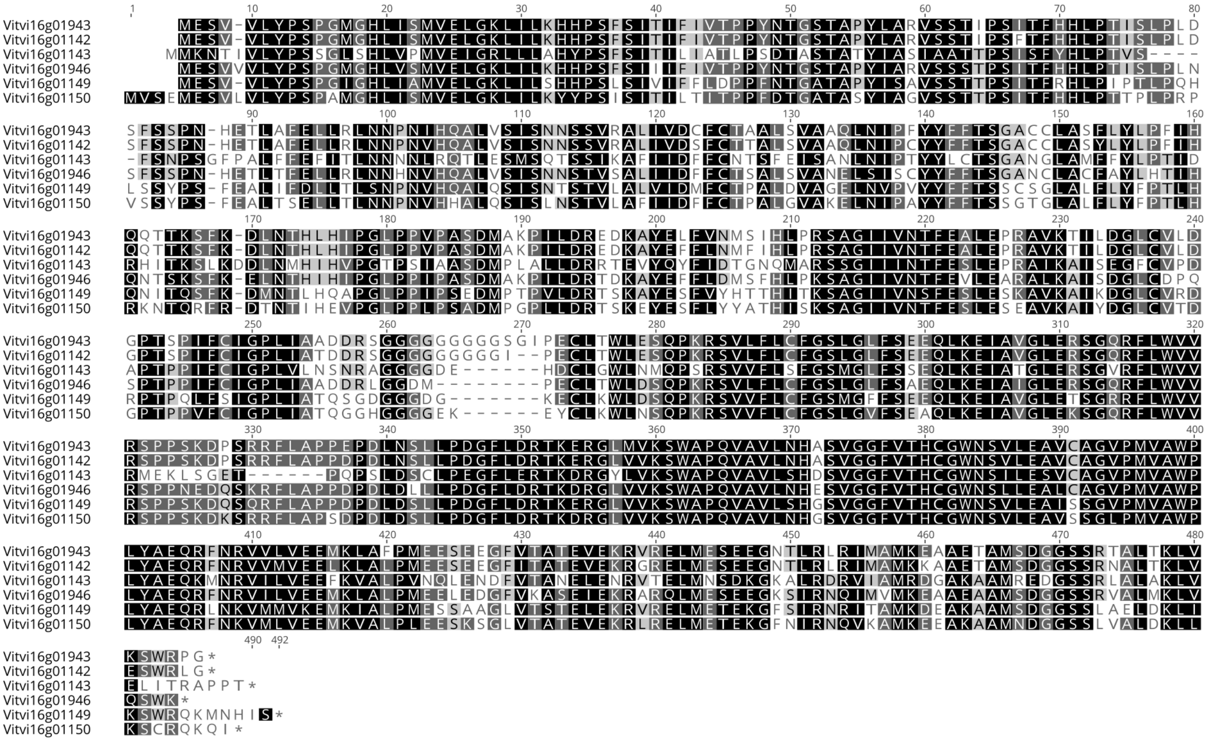

**Figure S4.** Alignment of the amino acid sequences of the six putative UGTs based on Ri genome sequence. The black and gray shading represents identical and similar amino acids, respectively.

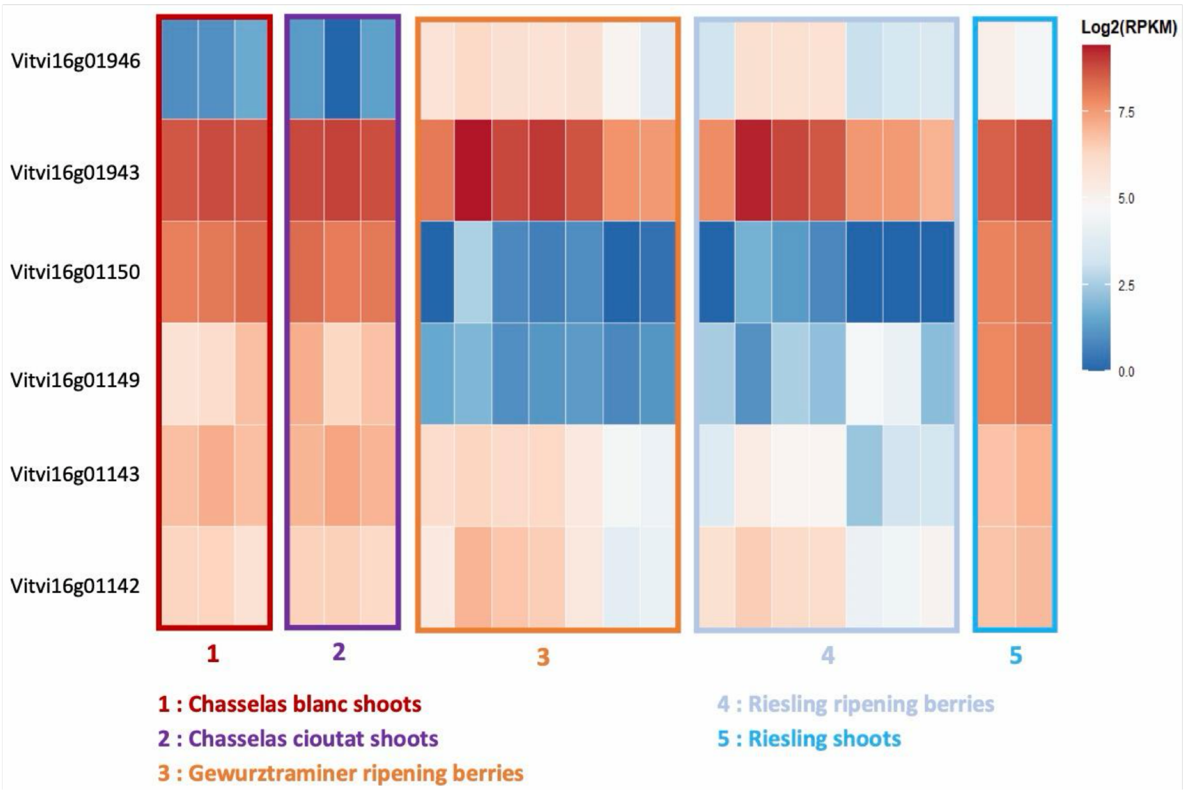

**Figure S5.** Overview of expression profiles of the six candidate UGTs in various tissues from different grape varieties, based on publicly available RNA seq analyses. RPKM = Reads per kilo base per million mapped reads.

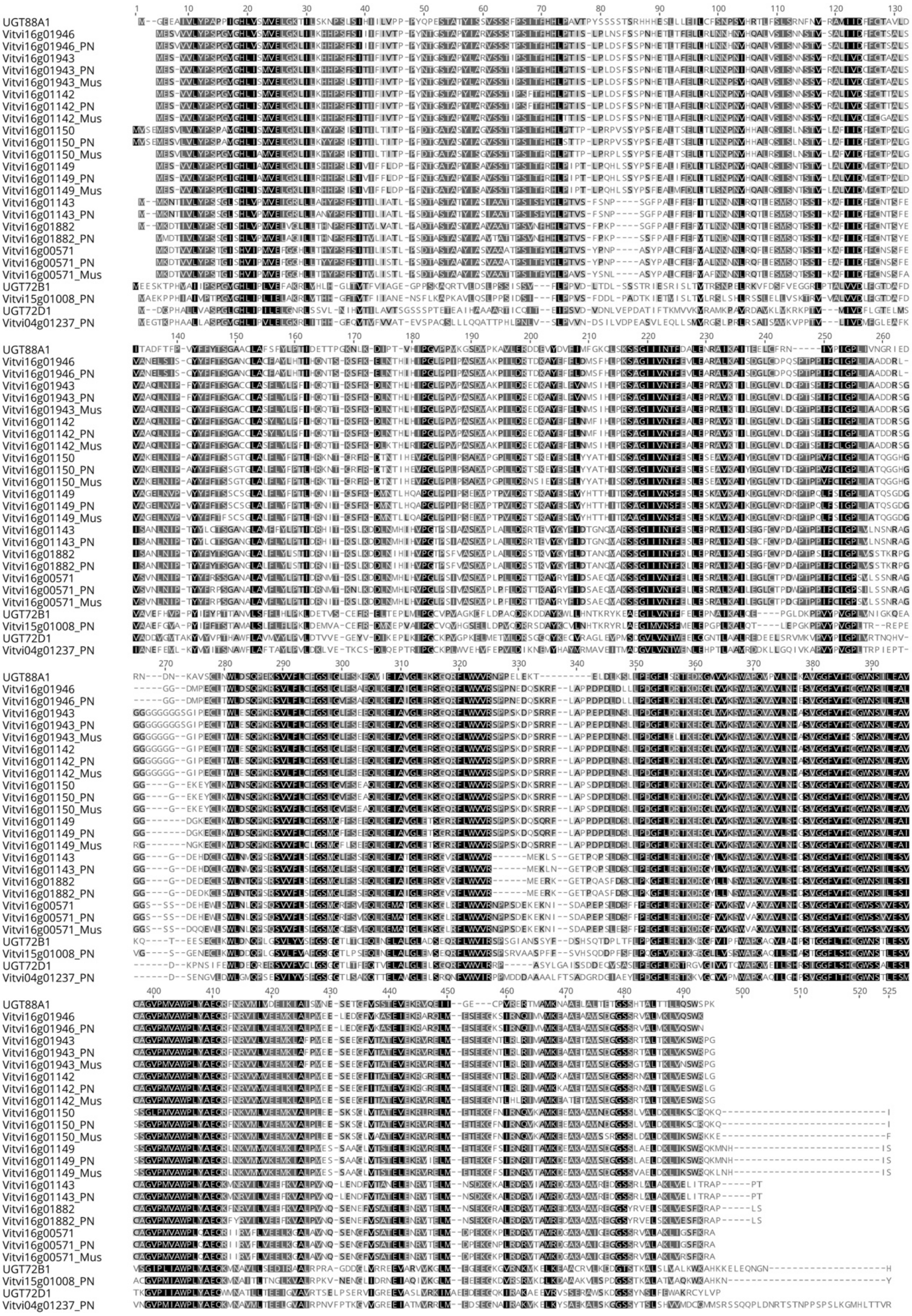

**Figure S6.** Alignment of the amino acid sequences of selected UGTs from *A. thaliana*, *V. vinifera* cv Riesling, *V. vinifera* PN40024 and *Muscadinia rotundifolia*.

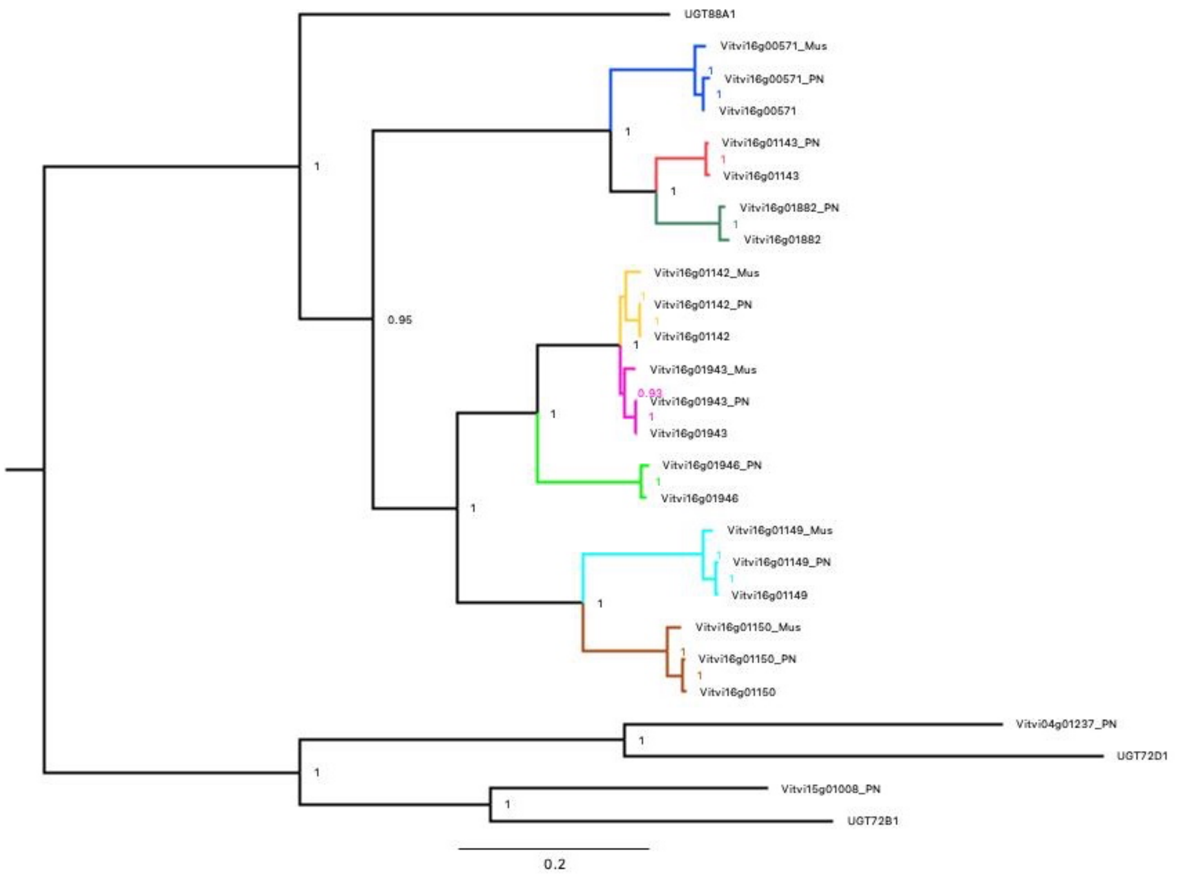

**Figure S7:** Bayesian phylogenetic tree built to estimate omega for the coding sequence of each protein.

<table>
<tr><th>Model</th><th colspan="2">dN/dS</th><th>p-value</th><th>Sites under positive selection</th></tr>
<tr><td colspan="5">Branch model</td></tr>
<tr><td rowspan="7">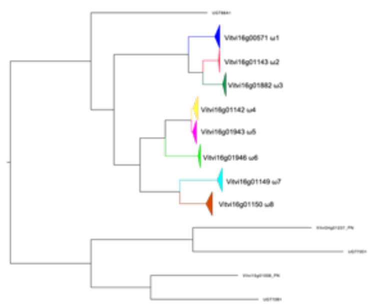</td><td>$\omega_0$=0.1705</td><td>-</td><td rowspan="7">8.57e-09</td><td rowspan="7">NA</td></tr>
<tr><td>$\omega_1$=0.6876</td><td>373-374</td></tr>
<tr><td>$\omega_2$=0.4492</td><td>373</td></tr>
<tr><td>$\omega_3$=0.2997</td><td>370-372</td></tr>
<tr><td>$\omega_4$=0.3386</td><td>397-400</td></tr>
<tr><td>$\omega_5$=0.2452</td><td>399-402</td></tr>
<tr><td>$\omega_6$=0.1909</td><td>403</td></tr>
</table>

| | | | | |
|---|---|---|---|---|
| | ω7=0.2310 | 389 | | |
| | ω8=0.3919 | 388-389 | | |
| **Branch-site model** | | | | |
| Vitvi16g01943+Vitvi16g01142 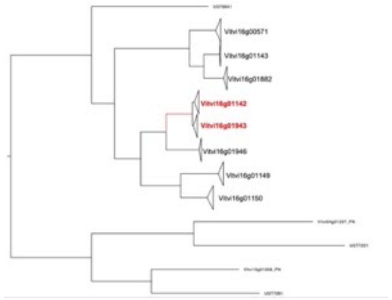 | **ω0= 0.183**<br>**ω1= 0.183** | p=0.782 | 0.023 | 161 (0.846)<br>462 (0.749) |
| | **ω0=1**<br>**ω1=1** | p=0.212 | | |
| | **ω0= 0.183**<br>ω1=437.316 | p=0.005 | | |
| | **ω0=1**<br>ω1=437.316 | p=0.001 | | |
| Vitvi16g01150 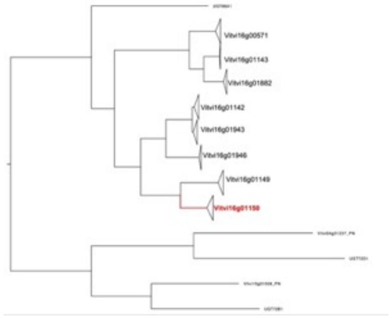 | **ω0=0. 184**<br>**ω1=0.184** | p=0.788 | 0.039 | 0 |
| | **ω0=1**<br>**ω1=1** | p=0.205 | | |
| | **ω0=0.184**<br>ω1=999 | p=0.006 | | |
| | **ω0=1**<br>**ω1=999** | p=0.002 | | |
| MxS2 | | | | |
| Vitvi16g01943+Vitvi16g01142 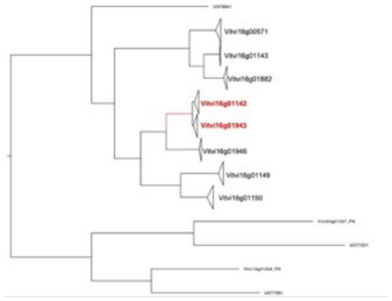 | **ω1=0.001** | p= 0.680 | 0.002 | 65 |
| | ω2=1.000 | p=0.295 | | |
| | **ω3=2.481** | p=0.025 | | |

**Figure S8.** Results of codeml (branch model and branch-site model) and Fitmodel (MxS2) analyses investigating selection pressures in different branches. The first column indicates the model tested, specifying the branches for which we detected positive (codeml) or relaxed selection (fitmodel). The second column shows the ω estimations for the background ($\omega_0$) and highlighted branches in codeml, and for the three ω categories in Fitmodel. The number of sites under purifying selection estimated from Fitmodel for the 8 UGT genes is shown next to codeml branch model omega values. The proportion of sites corresponding to the specified ω values/categories are shown for branch-site and MxS2 models. The third column indicates the p-value obtained from LRT comparing the model to the null hypothesis. The fourth column lists sites identified as being under positive selection with a posterior probability >0.7, where applicable.

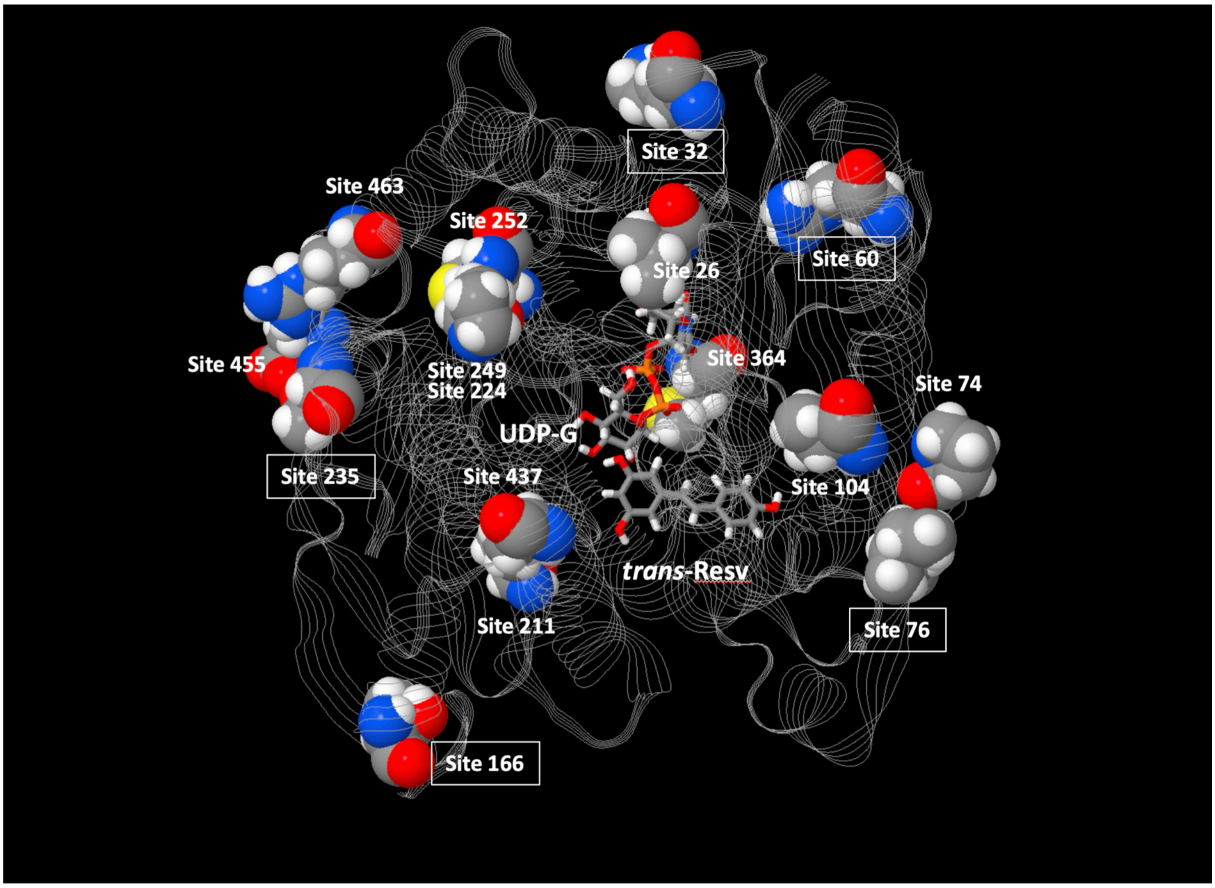

**Figure S9**. Positions of amino acid sites highlighted by the Pelican analysis located at the periphery of the UGT enzymes. Sites are illustrated in the Vitvi16g01943 structural model with *trans*-resveratrol (*trans*-Resv) and UDP-glucose (UDP-G) substrates. Sites discussed in the text are framed.

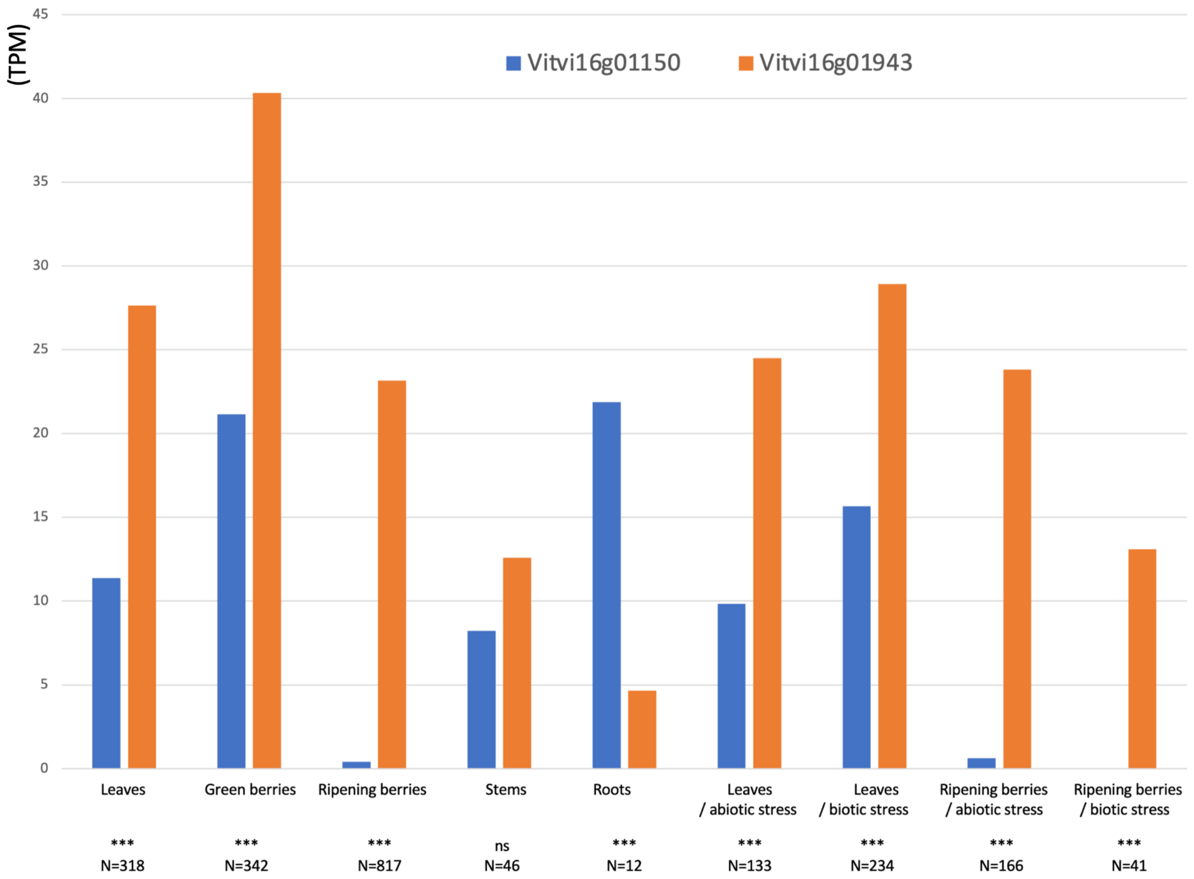

**Figure S10**. Expression profiles of the Vitivi16g01150 and Vitivi16g01943 genes in different grapevine organs. Expression of the Vitivi16g01150 and Vitivi16g01943 genes were quantified (in Transcripts Per Kilobase Million, TPM) in 2109 RNA-seq samples publicly available in the GREAT database (https://great.colmar.inrae.fr/). These RNA-seq samples include different grapevine organs, some of them submitted to abiotic or biotic stress as indicated. Numbers of RNA-seq samples and significant expression differences are indicated for each type of sample.

**Table S1**. Reproducible mQTLs based on non-targeted metabolomic analyses of canes and wood from the Ri x Gw population for the years 2020, 2021 and 2022.

| Identifier | Formula | Putative annotation | Tissues | Year | Map | Chr | LOD score | Peak LOD (cM) | LOD interval (cM) | Physical position of LOD interval (Mbps) |
|---|---|---|---|---|---|---|---|---|---|---|
| M345T195 | C16H26O8 | Picrocrocinic acid | Trunk | 2022 | GW | 5 | 19,2 | 15,08 | 15.1 - 17.5 | 3.51 - 4.05 |
| | | | Cane | 2020 | | | 11 | 16,67 | 15.1 - 19.1 | 3.51 - 4.44 |
| | | | Cane | 2021 | | | 11,8 | 16,67 | 15.1 - 19.1 | 3.51 - 4.44 |
| | | | Cane | 2022 | | | 9 | 15,88 | 13.5 - 17.5 | 3.44 - 4.05 |
| M493T380 | C22H38O12 | Acuminoside + formic acid | Trunk | 2022 | GW | 5 | 8,5 | 17,46 | 12.3 - 19.1 | 3 - 4.44 |
| | | | Cane | 2020 | | | 7 | 16,67 | 9.5 - 19.1 | 1.91 - 4.44 |
| | | | Cane | 2021 | | | 8,4 | 16,67 | 9.5 - 39.4 | 1.91 - 6.6 |
| | | | Cane | 2022 | | | 11,9 | 15,88 | 15.1 - 17.5 | 3.51 - 4.05 |
| M480T385 | C21H37NO11 | Unknown | Trunk | 2022 | GW | 5 | 16,4 | 15,88 | 15.1 - 17.5 | 3.53 - 4.05 |
| | | | Cane | 2020 | | | 7,5 | 19,85 | 9.5 - 22.6 | 1.91 - 5.52 |
| | | | Cane | 2021 | | | 13,8 | 19,85 | 9.5 - 19.8 | 1.91 - 4.80 |
| | | | Cane | 2022 | | | 10,7 | 15,08 | 15.1 - 17.5 | 3.53 - 4.05 |
| M485T385 | Unknown | Unknown | Trunk | 2022 | GW | 5 | 16,6 | 15,88 | 13.9 - 16.7 | 3.36 - 4.03 |
| | | | Cane | 2020 | | | 10,2 | 16,67 | 15.1 - 19.8 | 3.51 - 4.8 |
| | | | Cane | 2021 | | | 15,25 | 19,85 | 9.1 - 19.8 | 2.26 - 4.8 |
| | | | Cane | 2022 | | | 8,93 | 15,08 | 10.3 - 17.5 | 2.77 - 4.05 |
| M257T91 | C13H20O5 | Unknown | Trunk | 2022 | RI | 6 | 15,7 | 29,37 | 27 - 31 | 6.75 - 8.06 |
| | | | Cane | 2020 | | | 6,3 | 31,75 | 29.4 - 44.8 | 7.44 - 17.34 |

| | | | | | | | | | | |
|---|---|---|---|---|---|---|---|---|---|---|
| | | | Cane | 2021 | | | 6,3 | 37,31 | 32.5 - 49.2 | 8.58 - 18.59 |
| | | | Cane | 2022 | | | 6,4 | 31,36 | 27.4 - 33.7 | 6.78 - 9.06 |
| M589T535 | C30H32O10 | Pierotin A | Trunk | 2022 | RI | 6 | 9,5 | 31,36 | 28.2 - 40.1 | 7.14 - 14.03 |
| | | | Cane | 2020 | | | 3,2 | 29,37 | 18.3 - 43.3 | 4.57 - 17.31 |
| | | | Cane | 2021 | | | 12,1 | 39,29 | 31.4 - 46 | 8.1 - 17.68 |
| | | | Cane | 2022 | | | 6 | 29,77 | 25.8 - 35.3 | 6.28 - 10.73 |
| M699T530 | C39H38O12 | Dadahol A | Trunk | 2022 | RI | 6 | 5,7 | 29,77 | 22.6 - 42.5 | 4.57 - 16.64 |
| | | | Cane | 2020 | | | 29,4 | 31,75 | 18.3 - 43.3 | 4.57 - 17.31 |
| | | | Cane | 2021 | | | 8,1 | 38,9 | 15.9 - 43.3 | 3.86 - 17.31 |
| | | | Cane | 2022 | | | 3,9 | 31,36 | 25.8 - 35.3 | 6.28 - 10.73 |
| M463T92 | C20H32O12 | Unknown | Trunk | 2022 | RI | 6 | 14,4 | 31,36 | 28.6 - 33.7 | 7.28 - 9.06 |
| | | | Cane | 2020 | | | 8,3 | 37,71 | 28.6 - 44.5 | 7.28 - 17.45 |
| | | | Cane | 2021 | | | 4,5 | 36,52 | 31.4 - 43.3 | 8.1 - 17.31 |
| | | | Cane | 2022 | | | 4,8 | 31,36 | 18.3 - 33.7 | 4.57 - 9.06 |
| M525T116 | C25H34O12 | Unknown | Trunk | 2022 | RI | 6 | 10,1 | 31,36 | 27.8 - 35.3 | 7.07 - 10.55 |
| | | | Cane | 2020 | | | 5,5 | 29,37 | 22.2 - 43.3 | 5.27 - 17.31 |
| | | | Cane | 2021 | | | 3,2 | 36,91 | 23.4 - 43.3 | 5.56 - 17.31 |
| | | | Cane | 2022 | | | 3,2 | 36,91 | 23.4 - 43.3 | 5.56 - 17.31 |
| M729T534 | C40H42O13 | threo-carolignan E | Trunk | 2022 | RI | 6 | 8,6 | 39,29 | 31 - 47.2 | 8.07 - 17.94 |
| | | | Cane | 2020 | | | 3,1 | 25,01 | 0 - 46.4 | 0.8 - 17.75 |

| | | | | | | | | | | |
|---|---|---|---|---|---|---|---|---|---|---|
| | | | Cane | 2021 | | | 11,4 | 39,9 | 20.6 - 40.5 | 5.02 - 14.42 |
| | | | Cane | 2022 | | | 5,6 | 31,36 | 22.2 - 33.7 | 5.27 - 9.06 |
| M745T534 | C40H42O14 | Unknown | Trunk | 2022 | RI | 6 | 8,8 | 30,96 | 25.8 - 33.7 | 6.28 - 9.06 |
| | | | Cane | 2020 | | | 4,4 | 31,75 | 18.3 - 46.8 | 4.71 - 17.79 |
| | | | Cane | 2021 | | | 8,1 | 46,44 | 30.2 - 46.8 | 7.92 - 17.79 |
| | | | Cane | 2022 | | | 3 | 32,55 | 15.9 - 39.3 | 3.86 - 13.81 |
| M331T195 | C19H22O5 | Gibberellin A5/A7 | Trunk | 2022 | RI | 8 | 13,4 | 54,79 | 52.4 - 58.4 | 16.7 - 18.14 |
| | | | Cane | 2020 | | | 6,8 | 46,06 | 34.5 - 50.4 | 11.6 - 16.42 |
| | | | Cane | 2021 | | | 4,2 | 43,67 | 39.7 - 48.4 | 12.54 - 15.75 |
| | | | Cane | 2022 | | | 3,1 | 53,2 | 41.7 - 63.9 | 13.32 - 19.96 |
| M585T387 | C31H38O11 | Baccatin III | Trunk | 2022 | RI | 8 | 10,9 | 56,37 | 52.4 - 60.7 | 16.7 - 18.80 |
| | | | Cane | 2020 | | | 7,2 | 50,42 | 45.3 - 55.2 | 14.59 - 17.41 |
| | | | Cane | 2021 | | | 5,7 | 45,26 | 39.7 - 56 | 12.54 - 17.59 |
| | | | Cane | 2022 | | | 3,8 | 65,1 | 43.7 - 68.3 | 14.55 - 21.44 |
| M571T324 | C30H36O11 | Unknown | Trunk | 2022 | RI | 8 | 21,6 | 57,56 | 56 - 59.2 | 17.59 - 18.22 |
| | | | Cane | 2020 | | | 13 | 55,18 | 53.6 - 57.6 | 16.93 - 18.01 |
| | | | Cane | 2021 | | | 16 | 55,18 | 54.8 - 57.2 | 17.4 - 17.87 |
| | | | Cane | 2022 | | | 20 | 56,37 | 52.4 - 57.2 | 16.7 - 17.87 |
| M567T198 | C27H36O13 | Unknown | Trunk | 2022 | RI | 8 | 15,9 | 57,17 | 54.8 - 57.6 | 17.4 - 18.01 |
| | | | Cane | 2020 | | | 6,1 | 55,98 | 46.1 - 57.2 | 14.62 - 17.87 |

| | | | | | | | | | | |
|---|---|---|---|---|---|---|---|---|---|---|
| | | | Cane | 2021 | | | 5,2 | 45,26 | 39.7 - 50.4 | 12.54 - 16.42 |
| | | | Cane | 2022 | | | 4,7 | 53,2 | 43.7 - 59.9 | 14.55 - 18.67 |
| M325T97 | C15H20N2O6 | Unknown | Trunk | 2022 | RI | 10 | 6,3 | 8,73 | 6.7 - 12.7 | 2.37 - 5.53 |
| | | | Cane | 2020 | | | 5,6 | 10,32 | 0.8 - 18.7 | 0.25 - 8.03 |
| | | | Cane | 2021 | | | 13,7 | 10,32 | 6.7 - 12.7 | 2.37 - 5.53 |
| | | | Cane | 2022 | | | 12,8 | 8,73 | 6.7 - 10.7 | 2.37 - 4.56 |
| M325T97 | C15H20N2O6 | Unknown | Trunk | 2022 | GW | 10 | 7,66 | 8,73 | 2.4 - 9.9 | 1.63 - 4.54 |
| | | | Cane | 2020 | | | 6,05 | 7,54 | 5.6 - 9.9 | 2.43 - 4.54 |
| | | | Cane | 2021 | | | 3,18 | 7,94 | 0 - 9.9 | 1.05 - 4.54 |
| | | | Cane | 2022 | | | 3,66 | 5,16 | 0 - 18.3 | 1.05 - 6.08 |
| M485T97 | C21H30N2O11 | Unknown | Trunk | 2022 | RI | 10 | 6,4 | 9,92 | 2 - 11.5 | 0.37 - 5.01 |
| | | | Cane | 2020 | | | 5,1 | 15,08 | 2 - 16.3 | 0.37 - 6.56 |
| | | | Cane | 2021 | | | 13,3 | 45,26 | 6.7 - 13.1 | 2.37 - 5.92 |
| | | | Cane | 2022 | | | 12,1 | 8,73 | 4.8 - 9.1 | 1.58 - 3.82 |
| M247T113 | C6H15O8P | Glycerophosphoglycerol | Trunk | 2022 | RI | 14 | 3,9 | 78,2 | 64.3 - 79.4 | 25.93 - 30.75 |
| | | | Cane | 2020 | | | 4,4 | 52,39 | 31 - 61.9 | 13.27 - 25.8 |
| | | | Cane | 2021 | | | 6,2 | 64,31 | 52.4 - 79.4 | 22.41 - 30.75 |
| | | | Cane | 2022 | | | 4,5 | 65,5 | 46.8 - 67.5 | 21.95 - 26.72 |
| M423T97 | C18H32O11 | Unknown | Trunk | 2022 | RI | 14 | 4,2 | 67,48 | 59.9 - 73.8 | 25.49 - 29.57 |
| | | | Cane | 2020 | | | 12 | 64,7 | 64.3 - 66.3 | 25.93 - 26.57 |

| | | | | | | | | | | |
|---|---|---|---|---|---|---|---|---|---|---|
| | | | Cane | 2021 | | | 10,4 | 67,48 | 66.3 - 69.9 | 26.57 - 27.74 |
| | | | Cane | 2022 | | | 17,9 | 66,29 | 61.9 - 66.7 | 25.7 - 26.61 |
| M449T192 | C29H34O11 | Unknown | Trunk | 2022 | RI | 14 | 4,8 | 70,26 | 59.1 - 73.8 | 25.47 - 29.57 |
| | | | Cane | 2020 | | | 6,7 | 65,1 | 64.3 - 66.7 | 25.93 - 26.61 |
| | | | Cane | 2021 | | | 4,6 | 68,67 | 61.5 - 79.4 | 25.67 - 30.75 |
| | | | Cane | 2022 | | | 8,9 | 66,29 | 56 - 66.7 | 24.06 - 26.61 |
| M373T66 | C16H22O10 | Scologanate | Trunk | 2022 | GW | 15 | 13,5 | 28,98 | 26.6 - 34.9 | 16 - 18.17 |
| | | | Cane | 2020 | | | 6,2 | 27 | 23 - 29.4 | 14.85 - 19.9 |
| | | | Cane | 2021 | | | 5,4 | 28,89 | 26.6 - 29.4 | 16 - 20.04 |
| | | | Cane | 2022 | | | 7,4 | 28,89 | 26.6 - 42.1 | 16 - 20.1 |
| M197T93 | C10H12O4 | Cantharidine | Trunk | 2022 | GW | 15 | 7,8 | 24,61 | 19.1 - 29.4 | 13.75 - 16.37 |
| | | | Cane | 2020 | | | 7,9 | 26,99 | 23 - 28.6 | 14.85 - 16.28 |
| | | | Cane | 2021 | | | 8,7 | 27,78 | 25 - 28.6 | 15.8 - 16.28 |
| | | | Cane | 2022 | | | 7,5 | 24,61 | 23 - 32.55 | 14.85 - 16.6 |
| M227T94 | C11H14O5 | Genipin/ 3,4,5 Trimethoxyphenylacetic acid | Trunk | 2022 | GW | 15 | 4,5 | 25,01 | 21 - 38.5 | 14.47 - 19.76 |
| | | | Cane | 2020 | | | 4,7 | 26,99 | 19.1 - 29.4 | 13.75 - 16.37 |
| | | | Cane | 2021 | | | 3,3 | 28,18 | 23 - 38.5 | 14.85 - 19.76 |
| | | | Cane | 2022 | | | 5,2 | 26,59 | 24.6 - 42.1 | 15.7 - 20.1 |
| M389T225 | C20H22O8 | Piceid | Trunk | 2022 | RI | 16 | 16,8 | 44,47 | 42.5 - 45.3 | 18.28 - 19.82 |
| | | | Cane | 2020 | | | 17,4 | 44,47 | 42.5 - 46.1 | 18.28 - 19.9 |

| | | | | | | | | | | |
|---|---|---|---|---|---|---|---|---|---|---|
| | | | Cane | 2021 | | | 13,6 | 44,47 | 35.3 - 43.9 | 16.49 - 20.04 |
| | | | Cane | 2022 | | | 15,9 | 44,47 | 42.1 - 48.8 | 18.13 - 21.56 |
| M227T225 | C14H12O3 | Resveratrol | Trunk | 2022 | RI | 16 | 17,2 | 44,47 | 42.1 - 44.9 | 18.13 - 19.82 |
| | | | Cane | 2020 | | | 16,3 | 44,47 | 42.5 - 46.1 | 18.28 - 19.9 |
| | | | Cane | 2021 | | | 13,4 | 44,47 | 42.5 - 47.2 | 18.28 - 20.05 |
| | | | Cane | 2022 | | | 16,8 | 44,47 | 42.1 - 47.6 | 18.13 - 20.28 |
| M567T199 | C27H36O13 | lariciresinol 4-O-glucoside + Formic acid | Trunk | 2022 | RI | 17 | 5,9 | 47,71 | 43.7 - 62.8 | 1.65 - 5.39 |
| | | | Cane | 2020 | | | 14,2 | 52,87 | 52.87 - 54.1 | 11.25 - 11.58 |
| | | | Cane | 2021 | | | 8,7 | 52,87 | 45.3 - 54.1 | 8.51 - 11.58 |
| | | | Cane | 2022 | | | 13,1 | 45,3 | 45.3 - 51.7 | 8.51 - 11 |
| M435T334 | C21H24O10 | Phloridzin | Trunk | 2022 | GW | 18 | 14,3 | 67,48 | 60.3 - 71.8 | 14.2 - 23.5 |
| | | | Cane | 2020 | | | 14,1 | 71,85 | 70.3 - 75.4 | 22.83 - 26.46 |
| | | | Cane | 2021 | | | 9,1 | 71,85 | 70.3 - 71.8 | 22.83 - 23.5 |
| | | | Cane | 2022 | | | 16,4 | 74,63 | 71.5 - 75.4 | 23.26 - 26.46 |
| M435T98 | C20H36O10 | Ellagic acid xylopyranoside | Trunk | 2022 | GW | 18 | 64,4 | 36,1 | 35.7 - 36.1 | 7.89 - 8.16 |
| | | | Cane | 2020 | | | 9,9 | 37,31 | 33.3 - 46.4 | 7.52 - 10.1 |
| | | | Cane | 2021 | | | 9,3 | 36,51 | 32.1 - 41.7 | 7.31 - 9.42 |
| | | | Cane | 2022 | | | 19,6 | 36,12 | 34.9 - 36.5 | 7.67 - 8.25 |
| M433T136 | C20H34O10 | Unknown | Trunk | 2022 | GW | 18 | 23,9 | 35,72 | 34.9 - 36.1 | 7.67 - 8.16 |
| | | | Cane | 2020 | | | 9,2 | 36,51 | 33.3 - 36.5 | 7.52 - 8.25 |

| | | | | | | | | | | |
|---|---|---|---|---|---|---|---|---|---|---|
| | | | Cane | 2021 | | | 8,4 | 34,93 | 33.3 - 46.4 | 7.52 - 10.1 |
| | | | Cane | 2022 | | | 13 | 36,12 | 35.7 - 36.5 | 7.89 - 8.25 |
| M329T95 | C14H18O9 | 1-O-vanilloyl-beta-D-glucose | Trunk | 2022 | GW | 18 | 3,7 | 78,2 | 60.3 - 90.1 | 14.2 - 34.84 |
| | | | Cane | 2020 | | | 8 | 77,01 | 74.2 - 78.2 | 23.79 - 29.92 |
| | | | Cane | 2021 | | | 3,8 | 78,99 | 76.2 - 89.7 | 26.95 - 34.64 |
| | | | Cane | 2022 | | | 4,9 | 90,1 | 76.2 - 90.1 | 26.95 - 34.84 |
| M329T116 | C14H18O9 | 1-O-vanilloyl-beta-D-glucose | Trunk | 2022 | GW | 18 | 32,6 | 78,2 | 77 - 78.2 | 27.67 - 29.92 |
| | | | Cane | 2020 | | | 3,1 | 82,57 | 22.6 - 85.7 | 4.42 - 33.79 |
| | | | Cane | 2021 | | | 4,8 | 71,85 | 70.3 - 82.2 | 22.83 - 31.52 |
| | | | Cane | 2022 | | | 6,6 | 77 | 71.5 - 80.2 | 23.26 - 31.52 |
| M525T97 | C24H30O13 | Inumakilactone A glycoside | Trunk | 2022 | GW | 18 | 9,8 | 4,76 | 0 - 6.4 | 0.34 - 1.41 |
| | | | Cane | 2020 | | | 5,8 | 5,95 | 0 - 12.3 | 0.34 - 2.19 |
| | | | Cane | 2021 | | | 4,9 | 2,78 | 0 - 20.6 | 0.34 - 4.2 |
| | | | Cane | 2022 | | | 5,3 | 2,78 | 0 - 24.2 | 0.34 - 4.62 |
| M435T334 | C21H24O10 | Phloridzin | Trunk | 2022 | RI | 18 | 12,1 | 79,04 | 78.6 - 80.6 | 29.51 - 30.54 |
| | | | Cane | 2020 | | | 3,1 | 78,65 | 48.9 - 90.2 | 12.18 - 34.69 |
| | | | Cane | 2021 | | | 3,5 | 84,2 | 63.6 - 90.6 | 19.84 - 34.77 |
| | | | Cane | 2022 | | | 4,9 | 67,92 | 54 - 74.7 | 13.61 - 28.36 |
| M525T97 | C24H30O13 | Inumakilactone A glycoside | Trunk | 2022 | RI | 18 | 15,2 | 4,76 | 0 - 4.8 | 0 - 1.64 |
| | | | Cane | 2020 | | | 7,6 | 4,76 | 0 - 7.9 | 0 - 2.32 |

| | | | | | | | | | | |
|---|---|---|---|---|---|---|---|---|---|---|
| | | | Cane | 2021 | | | 5,3 | 1,19 | 0 - 14.3 | 0 - 4.05 |
| | | | Cane | 2022 | | | 5,5 | 0 | 0 - 6 | 0 - 1.78 |
| M471T286 | C28H22O7 | Scirpusin A | Trunk | 2022 | RI | 18 | 9,1 | 71,1 | 68.3 - 74.7 | 22.97 - 28.36 |
| | | | Cane | 2020 | | | 3,1 | 71,1 | 65.9 - 91 | 22.24 - 34.89 |
| | | | Cane | 2021 | | | 5,9 | 72,68 | 65.5 - 88.2 | 21.66 - 33.92 |
| | | | Cane | 2022 | | | 11,7 | 71,1 | 68.3 - 74.7 | 22.97 - 28.36 |
| M429T300 | C18H22O12 | Unknown | Trunk | 2022 | RI | 19 | 5,8 | 5,96 | 0 - 11.1 | 1.65 - 5.39 |
| | | | Cane | 2020 | | | 5 | 0,4 | 0 - 8.7 | 1.65 - 3.98 |
| | | | Cane | 2021 | | | 3,7 | 0,4 | 0 - 8.7 | 1.65 - 3.98 |
| | | | Cane | 2022 | | | 3,1 | 2,78 | 0 - 30.2 | 2.34 - 9.97 |

**Table S2.** Identifier and functional annotation of the genes located in the confidence interval of the piceid mQTL on chromosome 16 of Riesling

| PN40024.v4.2ID | V1_ID | Refseq_ID | Functional_annotation | Best_arabidopsis_match |
|---|---|---|---|---|
| Vitvi16g01163 | VIT_16s0050g01850 | LOC100257581 | AAA-type ATPase | AT5G52882 |
| Vitvi16g01957 | VIT_16s0050g01830 | LOC100252698 | AAA-type ATPase | AT5G52882 |
| Vitvi16g01144 | VIT_16s0050g01620 | LOC100245835 | ABC transporter G member 14 | AT5G52860 |
| Vitvi16g01979 | VIT_16s0050g02220 | LOC100233088 | Acidic endochitinase (CHIB1) | AT5G24090 |
| Vitvi16g01082 | VIT_16s0050g00870 | LOC100243942 | Acyltransferase | AT5G23940 |
| Vitvi16g01123 | VIT_16s0050g01380 | NA | Ankyrin repeat protein | NA |
| Vitvi16g01915 | VIT_16s0050g00930 | LOC100268149 | AnthoMATE1, anthocyanin permease AM1 | AT4G25640 |
| Vitvi16g01911 | VIT_16s0050g00900 | LOC100256121 | AnthoMATE3, anthocyanin permease AM3 | AT4G25640 |
| Vitvi16g01978 | VIT_16s0050g02200 | LOC100263797 | ARM repeat protein-like protein | NA |
| Vitvi16g01073 | VIT_16s0050g00780 | LOC100259320 | Arsenite transport protein (ArsB) | AT1G02260 |
| Vitvi16g01916 | VIT_16s0050g01010 | LOC100259304 | ATP binding protein | AT5G52545 |
| Vitvi16g01099 | VIT_16s0050g01090 | LOC100268009 | Beta-carotene hydroxylase | AT5G52570 |
| Vitvi16g01078 | VIT_16s0050g00830 | LOC100249245 | Beta-ketoacyl-CoA synthase | AT5G23940 |
| Vitvi16g01161 | VIT_16s0050g01790 | LOC100262972 | BTI3 (VIRB2- interacting protein 3) | AT2G46170 |
| Vitvi16g01080 | VIT_16s0050g00850 | LOC100249100 | C2 domain-containing protein | AT1G75350 |
| Vitvi16g01928 | VIT_16s0050g01300 | LOC100242355 | C2 domain-containing protein | AT5G23950 |
| Vitvi16g01929 | VIT_16s0050g01310 | LOC100852496 | C2 domain-containing protein | AT5G23950 |
| Vitvi16g01932 | VIT_16s0050g01400 | LOC104882216 | C2 domain-containing protein | AT3G16510 |
| Vitvi16g01166 | VIT_16s0050g01870 | LOC100254162 | CAAX amino terminal protease | AT5G60750 |
| Vitvi16g01197 | VIT_16s0050g02360 | LOC100246753 | Carrier protein CGI-69, Mitochondrial | AT4G27940 |
| Vitvi16g01973 | VIT_16s0050g02050 | NA | Casein kinase II subunit beta' | AT4G17640 |
| Vitvi16g01980 | VIT_16s0050g02230 | LOC100251796 | Chitinase [*Vitis vinifera*] | AT5G24090 |
| Vitvi16g01126 | VIT_16s0050g01420 | LOC100257712 | Coiled-coil domain-containing protein 111 | AT5G52800 |
| Vitvi16g01965 | VIT_16s0050g01960 | NA | Curculin (mannose-binding) lectin | AT5G24080 |
| Vitvi16g01964 | VIT_16s0050g01950 | LOC100247451 | Disease resistance family protein / LRR family | AT2G34930 |
| Vitvi16g01970 | VIT_16s0050g02010 | LOC100853108 | EIX receptor 1 | AT2G34930 |
| Vitvi16g01967 | VIT_16s0050g01980 | LOC100254307 | EIX receptor 2 | AT2G34930 |
| Vitvi16g01969 | GSVIVT00037814001 | LOC100256083 | EIX receptor 2 | AT2G34930 |
| Vitvi16g01076 | VIT_16s0050g00820 | LOC100254343 | EMB1691 (embryo defective 1691) | AT4G09980 |
| Vitvi16g01135 | VIT_16s0050g01500 | LOC100254325 | EMB2261 (embryo defective 2261) | AT5G52850 |
| Vitvi16g01162 | VIT_16s0050g01810 | LOC100257844 | F-box protein | AT5G52880 |
| Vitvi16g01090 | VIT_16s0050g00980 | LOC100252447 | ferric reduction oxidase 2 | AT1G01580 |
| Vitvi16g01091 | VIT_16s0050g00990 | LOC100247332 | ferric reduction oxidase 4 | AT5G23980 |
| Vitvi16g01102 | VIT_16s0050g01140 | LOC104877401 | Glutamine cyclotransferase | AT4G25720 |
| Vitvi16g01938 | VIT_16s0050g01530 | LOC100266393 | HcrVf2 protein | AT2G34930 |
| Vitvi16g01940 | VIT_16s0050g01550 | NA | HcrVf2 protein | AT2G34930 |
| Vitvi16g01103 | VIT_16s0050g01150 | LOC100257736 | Heat shock protein 90-1 | AT5G52640 |

| | | | | |
|---|---|---|---|---|
| Vitvi16g01926 | VIT_16s0050g01240 | LOC100252599 | Heavy-metal-associated domain-containing protein | AT1G01490 |
| Vitvi16g01074 | VIT_16s0050g00790 | LOC100264629 | Hydroxyproline-rich glycoprotein | AT4G25620 |
| Vitvi16g01199 | VIT_16s0050g02370 | LOC100258742 | Inorganic phosphate transporter 1-3 | AT5G43360 |
| Vitvi16g01175 | VIT_16s0050g02090 | NA | Kinesin motor | NA |
| Vitvi16g01914 | VIT_16s0050g00920 | LOC104882194 | MATE efflux family protein | AT4G00350 |
| Vitvi16g01915 | VIT_16s0050g00910 | LOC100250967 | MATE efflux family protein | AT4G25640 |
| Vitvi16g01923 | VIT_16s0050g01190 | LOC100247478 | Metal ion binding | AT1G01490 |
| Vitvi16g01113 | VIT_16s0050g01290 | LOC100256110 | Metal transporter CNNM4 (Cyclin-M4) | AT5G52790 |
| Vitvi16g01097 | VIT_16s0050g01070 | LOC100245692 | MIND (accumulation and replication of chloroplast 11) | AT5G24020 |
| Vitvi16g01188 | VIT_16s0050g02280 | LOC100260363 | Multidrug resistance-associated protein 14 | AT3G59140 |
| Vitvi16g01106 | VIT_16s0050g01180 | LOC100252586 | Myb family | AT5G52660 |
| Vitvi16g01108 | NA | LOC100854849 | Unknown | NA |
| Vitvi16g01117 | NA | LOC104882189 | NF-kappa-B inhibitor cactus-like isoform X1 | NA |
| Vitvi16g01141 | NA | LOC100256100 | Leucine-rich repeat receptor-like protein kinase PXL1 | NA |
| Vitvi16g01151 | NA | NA | Late embryogenesis abundant protein | NA |
| Vitvi16g01159 | NA | LOC104882207 | Late embryogenesis abundant protein | NA |
| Vitvi16g01198 | NA | LOC104882171 | Mavicyanin-like | NA |
| Vitvi16g01908 | NA | LOC100264493 | short-chain dehydrogenase TIC 32, chloroplastic | NA |
| Vitvi16g01910 | NA | LOC104882220 | B3 domain-containing protein | NA |
| Vitvi16g01921 | NA | LOC100255958 | probable polygalacturonase | NA |
| Vitvi16g01922 | NA | LOC100250814 | probable polygalacturonase | NA |
| Vitvi16g01930 | NA | LOC104882218 | Unknown | NA |
| Vitvi16g01935 | NA | LOC104882214 | Protein SRC2 | NA |
| Vitvi16g01936 | NA | LOC104882187 | Protein SRC2 | NA |
| Vitvi16g01937 | NA | LOC100249228 | LRR receptor-like serine/threonine-protein kinase FLS2 | NA |
| Vitvi16g01948 | NA | LOC104882210 | Unknown | NA |
| Vitvi16g01953 | NA | LOC104882182 | Late embryogenesis abundant protein | NA |
| Vitvi16g01959 | NA | LOC100252455 | Putative membrane-associated kinase regulator 6 | NA |
| Vitvi16g01134 | VIT_16s0050g01490 | LOC100249252 | NADH dehydrogenase (ubiquinone) 1 alpha subcomplex 5 | AT5G52840 |
| Vitvi16g01191 | VIT_16s0050g02310 | LOC100263802 | Arabinogalactan peptide 20 | NA |
| Vitvi16g01912 | GSVIVT01028884001 | LOC104882193 | Protein detoxification 35 | NA |
| Vitvi16g01924 | VIT_16s0050g01210 | NA | Unknown | NA |
| Vitvi16g01152 | GSVIVT00037782001 | LOC100250793 | Nodulin MtN21 family | AT5G64700 |

| | | | | |
|---|---|---|---|---|
| Vitvi16g01958 | VIT_16s0050g01840 | LOC100247304 | Nucleoside triphosphatase | AT5G52882 |
| Vitvi16g01127 | VIT_16s0050g01430 | LOC100259461 | Ornithine cyclodeaminase | AT5G52810 |
| Vitvi16g01184 | VIT_16s0050g02250 | LOC100248393 | Pentatricopeptide (PPR) repeat | AT3G49140 |
| Vitvi16g01129 | VIT_16s0050g01450 | LOC100247467 | Pentatricopeptide (PPR) repeat-containing protein | AT4G21190 |
| Vitvi16g01173 | VIT_16s0050g02070 | NA | Pentatricopeptide repeat-containing protein | NA |
| Vitvi16g01100 | VIT_16s0050g01120 | NA | Polygalacturonase GH28 | AT3G61490 |
| Vitvi16g01920 | VIT_16s0050g01110 | LOC100262858 | Polygalacturonase GH28 | AT3G48950 |
| Vitvi16g01087 | VIT_16s0050g00960 | LOC100262997 | Prolyl-tRNA synthetase | AT5G52520 |
| Vitvi16g01094 | VIT_16s0050g01020 | LOC100254190 | Protein kinase family | AT5G24010 |
| Vitvi16g01182 | VIT_16s0050g02180 | LOC100262029 | Pyruvate kinase isozyme G, chloroplast precursor | AT5G52920 |
| Vitvi16g01167 | VIT_16s0050g01880 | LOC100243904 | Receptor kinase homolog LRK10 | AT5G24080 |
| Vitvi16g04412 | VIT_16s0050g02060 | LOC100250224 | Receptor kinase homolog LRK10 | AT1G66920 |
| Vitvi16g01192 | VIT_16s0050g02320 | LOC100244982 | Regulator of nonsense transcripts 1 | AT1G65810 |
| Vitvi16g01193 | VIT_16s0050g02330 | LOC100267290 | Regulator of nonsense transcripts 1 | AT1G65810 |
| Vitvi16g01194 | VIT_16s0050g02340 | LOC104882173 | Regulator of nonsense transcripts 1 | AT1G65810 |
| Vitvi16g01195 | VIT_16s0050g02350 | LOC100257019 | Regulator of nonsense transcripts 1 | AT1G65810 |
| Vitvi16g01118 | VIT_16s0050g01330 | LOC100252438 | Retrovirus Pol polyprotein from transposon TNT 1-94 | AT3G29785 |
| Vitvi16g01119 | VIT_16s0050g01340 | LOC100242184 | Retrovirus Pol polyprotein from transposon TNT 1-94 | AT3G29785 |
| Vitvi16g01081 | VIT_16s0050g00860 | LOC100261205 | Ribosomal protein L31 | NA |
| Vitvi16g01104 | VIT_16s0050g01160 | LOC100257720 | Ribosomal protein S10 (RPS10C) 40S | NA |
| Vitvi16g01083 | VIT_16s0050g00880 | LOC100266267 | RRNA 2'-O-methyltransferase fibrillarin | AT5G52470 |
| Vitvi16g01086 | VIT_16s0050g00950 | LOC100261057 | Scarecrow-like transcription factor 8 | AT5G52510 |
| Vitvi16g01909 | VIT_16s0050g00800 | LOC100254207 | Sec-independent protein translocase protein TatA | AT5G52440 |
| Vitvi16g01966 | VIT_16s0050g01970 | NA | Serine/threonine protein kinase LRK19 | NA |
| Vitvi16g01101 | VIT_16s0050g01130 | LOC100267993 | Serine/threonine-protein kinase | AT1G01540 |
| Vitvi16g01950 | VIT_16s0050g01740 | LOC100249217 | Serine-threonine protein kinase, plant-type | NA |
| Vitvi16g01136 | VIT_16s0050g01510 | LOC100244083 | SGT1-2 | AT4G11260 |
| Vitvi16g01907 | VIT_16s0050g00750 | LOC100242214 | Short-chain dehydrogenase/reductase | AT4G23420 |
| Vitvi16g01070 | VIT_16s0050g00760 | LOC100247484 | Short-chain dehydrogenase/reductase | AT4G23420 |
| Vitvi16g01164 | VIT_16s0050g01860 | LOC100242167 | SLAH3 (SLAC1 Homologue 3) | AT5G24030 |
| Vitvi16g01963 | VIT_16s0050g01940 | NA | S-locus lectin protein kinase | NA |
| Vitvi16g01931 | VIT_16s0050g01390 | LOC104882217 | SRC2/SRC2 (soybean gene regulated BY cold-2) | NA |
| Vitvi16g01942 | VIT_16s0050g01570 | LOC100266422 | Synbindin TRAPPC1 | AT1G51160 |
| Vitvi16g01181 | VIT_16s0050g02160 | LOC100246656 | Timeless ATIM | AT5G52910 |

| | | | | |
|---|---|---|---|---|
| Vitvi16g01130 | VIT_16s0050g01460 | LOC100254350 | Translocation protein SEC63 | AT1G79940 |
| Vitvi16g01075 | VIT_16s0050g00810 | LOC104882195 | Ubiquitin-protein ligase | NA |
| Vitvi16g01149 | VIT_16s0050g01660 | LOC100254171 | UDP-glucoronosyl/UDP-glucosyl transferase UGT88A1 | NA |
| Vitvi16g01142 | VIT_16s0050g01590 | LOC100256130 | UDP-glucose: anthocyanidin 5,3-O-glucosyltransferase | AT3G16520 |
| Vitvi16g01150 | VIT_16s0050g01680 | LOC100268157 | UDP-glucose: anthocyanidin 5,3-O-glucosyltransferase | AT3G16520 |
| Vitvi16g01943 | VIT_16s0050g01580 | LOC100261214 | UDP-glucose: anthocyanidin 5,3-O-glucosyltransferase | AT3G16520 |
| Vitvi16g01143 | VIT_16s0050g01600 | LOC100247310 | UDP-glycosyltransferase 88A4 | AT3G16520 |
| Vitvi16g01946 | VIT_16s0050g01610 | LOC100250976 | UDP-glycosyltransferase 88A4 | AT3G16520 |
| Vitvi16g01089 | VIT_16s0050g00970 | LOC104882192 | Unknown | NA |
| Vitvi16g01098 | VIT_16s0050g01080 | LOC100855090 | UDP-N-acetylmuramate--L-alanine ligase | NA |
| Vitvi16g01105 | VIT_16s0050g01170 | LOC104882190 | Unknown | NA |
| Vitvi16g01112 | VIT_16s0050g01260 | LOC100266405 | unknown | AT5G52780 |
| Vitvi16g01121 | VIT_16s0050g01350 | LOC104882188 | Unknown | NA |
| Vitvi16g01146 | VIT_16s0050g01650 | LOC104882184 | Late embryogenesis abundant protein | NA |
| Vitvi16g01183 | VIT_16s0050g02190 | LOC100241527 | Unknown | NA |
| Vitvi16g01941 | VIT_16s0050g01560 | LOC100262982 | trafficking protein particle complex subunit 1 isoform X1 | NA |
| Vitvi16g01951 | VIT_16s0050g01750 | LOC104882183 | Late embryogenesis abundant protein | NA |
| Vitvi16g01952 | VIT_16s0050g01760 | LOC104882208 | Late embryogenesis abundant protein | NA |
| Vitvi16g01954 | VIT_16s0050g01770 | LOC104882181 | Late embryogenesis abundant protein | NA |
| Vitvi16g01960 | VIT_16s0050g01890 | NA | Unknown | NA |
| Vitvi16g01968 | VIT_16s0050g01990 | LOC100854653 | Unknown | NA |
| Vitvi16g01982 | VIT_16s0050g02300 | NA | Unknown | NA |
| Vitvi16g01079 | VIT_16s0050g00840 | LOC100244074 | Unknown protein | AT3G49070 |
| Vitvi16g01093 | VIT_16s0050g01000 | LOC100264478 | membrane protein YjcL | AT5G52540 |
| Vitvi16g01095 | VIT_16s0050g01050 | LOC100255969 | deSI-like protein At4g17486 | AT4G25660 |
| Vitvi16g01096 | VIT_16s0050g01060 | LOC100250830 | Unknown protein | AT4G25670 |
| Vitvi16g01128 | VIT_16s0050g01440 | LOC100252576 | Unknown protein | AT3G49140 |
| Vitvi16g01131 | VIT_16s0050g01470 | LOC100242331 | Unknown protein | AT5G62650 |
| Vitvi16g01185 | VIT_16s0050g02260 | LOC100243267 | Unknown protein | AT5G52950 |
| Vitvi16g01187 | VIT_16s0050g02270 | LOC100241531 | Unknown protein | AT5G52960 |
| Vitvi16g01190 | VIT_16s0050g02290 | LOC100255280 | Unknown protein | AT5G53220 |
| Vitvi16g01981 | VIT_16s0050g02240 | LOC100246662 | protein CURVATURE THYLAKOID 1A | AT4G01150 |
| Vitvi16g01171 | VIT_16s0050g02000 | LOC100261168 | WD40 | NA |
| Vitvi16g01132 | VIT_16s0050g01480 | LOC104882213 | WRKY DNA-binding protein 27 | NA |
| Vitvi16g01133 | GSVIVT00037762001 | LOC104882212 | WRKY DNA-binding protein 27 | AT5G52830 |
| Vitvi16g01071 | VIT_16s0050g00770 | LOC100242348 | YDA (YODA) | AT1G63700 |
| Vitvi16g01919 | VIT_16s0050g01100 | NA | Zinc finger (C2H2 type) family | AT1G72050 |

**Table S3.** Protein identity between the six UGTs sequences based on MUSCLE alignment.

| | Vitvi16g01943 | Vitvi16g01142 | Vitvi16g01143 | Vitvi16g01946 | Vitvi16g01946 | Vitvi16g01946 |
|---|---|---|---|---|---|---|
| Vitvi16g01943 | **100%** | **95%** | **56%** | **81%** | **68%** | **69%** |
| Vitvi16g01142 | **95%** | **100%** | **56%** | **81%** | **68%** | **69%** |
| Vitvi16g01143 | **56%** | **56%** | **100%** | **57%** | **52%** | **54%** |
| Vitvi16g01946 | **81%** | **81%** | **57%** | **100%** | **68%** | **69%** |
| Vitvi16g01149 | **68%** | **68%** | **52%** | **68%** | **100%** | **77%** |
| Vitvi16g01150 | **69%** | **69%** | **54%** | **69%** | **77%** | **100%** |

**Table S4.** Kinetic parameters of recombinant VLRSgt, Vitvi16g01943*,* Vitvi16g01142, Vitvi16g01946 and Vitvi16g01150 with quercetin, kaempferol or resveratrol as substrates, and UDP-glucose as sugar donor (200 µM). N.A. = No significant activity. Data are presented as means ± the standard error of the mean (n = 3).

| Substrate | $K_m$ (µM) | $K_{cat}$ ($s^{-1}$) | $K_{cat}/K_m$ ($s^{-1}$ $mM^{-1}$) |
|---|---|---|---|
| VLRSgt | | | |
| Quercetin | 58.76 ± 12.4 | 0.29 ± 0.07 | 4.9 |
| Kaempferol | 65.1 ± 14.1 | 0.85 ± 0.19 | 13.05 |
| Resveratrol | 98.5 ± 19.1 | 0.17 ± 0.07 | 1.72 |
| Vitvi16g01946 | | | |
| Quercetin | 445.9 ± 92 | 1.98 ± 0.29 | 4.4 |
| Kaempferol | 640.9 ± 102 | 1.27 ± 0.17 | 1.99 |
| Resveratrol | 51.8 ± 17.3 | 0.22 ± 0.06 | 4.24 |
| Vitvi16g01943 | | | |
| Quercetin | 329.5 ± 78.2 | 0.21 ± 0.06 | 0.65 |
| Kaempferol | 229.7 ± 63.5 | 1.06 ± 0.12 | 4.6 |
| Resveratrol | 71.2 ± 21.1 | 0.72 ± 0.14 | 10.1 |
| Vitvi16g01142 | | | |
| Quercetin | N.A. | N.A. | N.A. |
| Kaempferol | N.A. | N.A. | N.A. |
| Resveratrol | 341.4 ± 78 | 0.69 ± 0.15 | 2 |
| Vitvi16g01150 | | | |
| Quercetin | N.A. | N.A. | N.A. |
| Kaempferol | N.A. | N.A. | N.A. |
| Resveratrol | 7.2 ± 2.1 | 2.67 ± 0.55 | 370 |

**Table S5.** Kd (µM) calculated from the molecular dynamics trajectories.

| | Vitvi16g01943 | Vitvi16g01946 | Vitvi16g01150 |
|---|---|---|---|
| trans-Resvératrol | 1.14 ± 0.22 | 1.11 ± 0.36 | 0.78 ± 0.14 |
| cis-Resvératrol | 0.86 ± 0.16 | 2.04 ± 0.31 | 2.17 ± 0.22 |

**Table S6.** Results from Pelican analysis, highlighting statistically significant differences in amino acid composition between protein lineages. The first column lists the protein lineages investigated; the second column indicates the position of the amino acid in the protein sequence; the third column shows the amino acids that are more prevalent in the investigated protein lineages and their frequency estimated from Pelican (frequencies < 0.01 were disregarded), next the associated p-values, indicating the statistical significance of the observed differences.

| Proteins | Sites | Amino acid(s) prevailing in the scrutinized protein(s) and frequency | P-value |
|---|---|---|---|
| **Vitvi16g01943+Vitvi16g01142+ Vitvi16g01150+Vitvi16g01146** | 2 | V (1.000) | 0.045948 |
| | 17 | M (1.000) | 0.049142 |
| | 22 | S (0.863), A (0.137) | 0.028322 |
| | 32 | K (0.988) | 0.020802 |
| | 60 | R (0.642), G (0.358) | 0.014149 |
| | 76 | I (0.688), T (0.311) | 0.006417 |
| | 166 | T (0.817), N (0.179) | 0.029651 |
| | 211 | F (0.750), S (0.249) | 0.016778 |
| | 463 | Q (0.952), R (0.047) | 0.030999 |
| **Vitvi16g01943+Vitvi16g01142+ Vitvi16g01150** | 2 | V (0.999) | 0.045948 |
| | 125 | C (0.402), G (0.351), S (0.247) | 0.036286 |
| | 235 | T (0.996) | 0.023617 |
| **Vitvi16g01943+Vitvi16g01142** | 125 | C (0.420), S (0.301), G (0.276) | 0.022950 |
| | 235 | T (1.000) | 0.003805 |
| | 357 | L (0.630), R (0.370) | 0.032489 |
| **Vitvi16g01146** | 160 | H (1.000) | 0.033913 |
| | 174 | E (1.000) | 0.037675 |
| | 446 | A (0.999) | 0.039643 |
| **Vitvi16g01150** | 16 | A (0.999) | 0.045847 |
| **Vitvi16g01943** | 104 | I (0.999) | 0.035527 |
| | 357 | L (1.000) | 0.010039 |
| | 364 | M (1.000) | 0.032281 |
| | 387 | S (1.000) | 0.012479 |
| **Vitvi16g00571** | 20 | V (1.000) | 0.004674 |
| | 147 | P (0.666), S (0.334) | 0.007661 |
| | 256 | S (0.999) | 0.037951 |
| | 350 | F (1.000) | 0.004585 |
| | 369 | V (1.000) | 0.004595 |

| | | | |
|---|---|---|---|
| | 370 | A (1.000) | 0.004862 |
| | 390 | S (0.997) | 0.032659 |
| | 393 | V (1.000) | 0.004674 |
| | 408 | Q (1.000) | 0.012381 |
| | 414 | I (0.998) | 0.028158 |
| **Vitvi16g01143** | 26 | I (0.997) | 0.035492 |
| | 455 | D (0.999) | 0.028578 |
| **Vitvi16g01882** | 27 | A (0.545), V (0.454) | 0.023736 |
| | 160 | S (0.999) | 0.019575 |
| | 365 | L (1.000) | 0.022585 |
| | 481 | Y (1.000) | 0.006959 |
| **Vitvi16g01149** | 150 | C (0.999) | 0.033045 |
| | 224 | S (0.999) | 0.036052 |
| | 249 | Q (0.997) | 0.017633 |
| | 252 | S (0.999) | 0.023806 |
| **Vitvi16g01142** | 74 | T (0.998) | 0.036731 |
| | 437 | I (0.999) | 0.017098 |
| | 461 | H (0.775), R (0.225) | 0.017269 |

**Table S7.** Samples used in this study. The table lists the species from which the sequences were derived, the corresponding variety, the proteins analyzed and the accession numbers of the sequences retrieved from publicly available databases. Sequences with asterisks (*) were used in downstream analyses with a smaller dataset.

| **Species** | **Variety** | **Protein** | **Accession number** |
|---|---|---|---|
| *A. thaliana* | NA | UGT72B1 | AT4G01070 |
| | | UGT72B2 | AT1G01390 |
| | | UGT72B3 | AT1G01420 |
| | | UGT72D1 | AT2G18570 |
| | | UGT72C1 | AT4G36770 |
| | | UGT72E1 | AT3G50740 |
| | | UGT72E2 | AT5G66690 |
| | | UGT72E3 | AT5G26310 |
| | | UGT88A1 | AT3G16520 |
| *V. vinifera* | PN40024 | Vitvi16g01943 | Vitvi16g01943 |
| | | Vitvi16g01142 | Vitvi16g01142 |
| | | Vitvi16g01946 | Vitvi16g01946 |
| | | Vitvi16g01150 | Vitvi16g01150 |
| | | Vitvi16g01149 | Vitvi16g01149 |
| | | Vitvi16g01882 | Vitvi16g01882 |
| | | Vitvi16g01143 | Vitvi16g01143 |
| | | Vitvi16g00571 | Vitvi16g00571 |
| | | Vitvi18g00926 | Vitvi18g00926 |
| | | Vitvi04g01237 | Vitvi04g01237 |

| | | | |
|---|---|---|---|
| | | Vitvi16g00576 | Vitvi16g00576 |
| | | Vitvi16g00849 | Vitvi16g00849 |
| | | Vitvi04g01233 | Vitvi04g01233 |
| | | Vitvi02g00110 | Vitvi02g00110 |
| | | Vitvi15g00597 | Vitvi15g00597 |
| | | Vitvi15g01008 | Vitvi15g01008 |
| | | Vitvi15g01623 | Vitvi15g01623 |
| | Riesling | Vitvi16g01943 | VITVvi_vRiesFPS24_v1.0_P63.ver1.0.g630370 |
| | | Vitvi16g01142 | VITVvi_vRiesFPS24_v1.0_P63.ver1.0.g630380 |
| | | Vitvi16g01946 | VITVvi_vRiesFPS24_v1.0_P4.ver1.0.g489470 |
| | | Vitvi16g01150 | VITVvi_vRiesFPS24_v1.0_P4.ver1.0.g489540 |
| | | Vitvi16g01149 | VITVvi_vRiesFPS24_v1.0_P4.ver1.0.g489510 |
| | | Vitvi16g01882 | VITVvi_vRiesFPS24_v1.0_P267.ver1.0.g374900 |
| | | Vitvi16g01143 | VITVvi_vRiesFPS24_v1.0_P4.ver1.0.g489460 |
| | | Vitvi16g00571 | VITVvi_vRiesFPS24_v1.0_P56.ver1.0.g625570 |
| | | Vitvi18g00926 | VITVvi_vRiesFPS24_v1.0_P401.ver1.0.g492000 |
| | | Vitvi04g01237 | VITVvi_vRiesFPS24_v1.0_P369.ver1.0.g451420 |
| | | Vitvi16g00576 | VITVvi_vRiesFPS24_v1.0_P56.ver1.0.g625580 |
| | | Vitvi16g00849 | VITVvi_vRiesFPS24_v1.0_P297.ver1.0.g392390 |
| | | Vitvi04g01233 | VITVvi_vRiesFPS24_v1.0_P369.ver1.0.g451380 |
| | | Vitvi02g00110 | VITVvi_vRiesFPS24_v1.0_P361.ver1.0.g444940 |
| | | Vitvi15g00597 | VITVvi_vRiesFPS24_v1.0_P333.ver1.0.g419860 |
| | | Vitvi15g01008 | VITVvi_vRiesFPS24_v1.0_H000059F_007.ver1.0.g101100 |
| | | Vitvi15g01623 | VITVvi_vRiesFPS24_v1.0_H000059F_007.ver1.0.g101120 |
| *M. rotundifolia* | Trayshed | Vitvi16g01943 | VITMroTrayshed_v2.0.hap1.chr16.ver2.0.g203780 |
| | | Vitvi16g01142 | VITMroTrayshed_v2.0.hap1.chr16.ver2.0.g203790 |
| | | Vitvi16g01150 | VITMroTrayshed_v2.0.hap1.chr16.ver2.0.g203870 |
| | | Vitvi16g01149 | VITMroTrayshed_v2.0.hap1.chr16.ver2.0.g203840 |
| | | Vitvi16g00571 | VITMroTrayshed_v2.0.hap1.chr16.ver2.0.g198220 |
| | | Vitvi18g00926 | VITMroTrayshed_v2.0.hap1.chr18.ver2.0.g226510 |
| | | Vitvi04g01237 | VITMroTrayshed_v2.0.hap1.chr04.ver2.0.g048680 |
| | | Vitvi04g01233 | VITMroTrayshed_v2.0.hap1.chr04.ver2.0.g048640 |
| | | Vitvi02g00110 | VITMroTrayshed_v2.0.hap1.chr02.ver2.0.g016020 |
| | | Vitvi15g00597 | VITMroTrayshed_v2.0.hap1.chr15.ver2.0.g187960 |
| | | Vitvi15g01008 | VITMroTrayshed_v2.0.hap1.chr15.ver2.0.g192070 |
